# Hierarchical determinants of the oxidation-induced mutational landscape in human cells

**DOI:** 10.1101/2024.04.08.588428

**Authors:** Cameron Cordero, Kavi P. M. Mehta, Tyler M. Weaver, Justin A. Ling, Bret D. Freudenthal, David Cortez, Steven A. Roberts

**Affiliations:** Department of Microbiology and Molecular Genetics, University of Vermont Cancer Center, University of Vermont, Burlington, VT 05405, USA; School of Molecular Biosciences and Center for Reproductive Biology, Washington State University, Pullman, WA 99164, USA; Department of Biochemistry, Vanderbilt University School of Medicine, Nashville, TN 37232, USA; Department of Comparative Biosciences, School of Veterinary Medicine, University of Wisconsin, Madison, WI 53706, USA; Department of Biochemistry and Molecular Biology, Department of Cancer Biology, University of Kansas Medical Center, Kansas City, KS 66160, USA; University of Kansas Cancer Center, Kansas City, KS 66160, USA

## Abstract

8-oxoguanine (8-oxoG) is a common oxidative DNA lesion, which causes G>T substitutions that compose COSMIC single base substitution signature 18 (SBS18) in human cancers. Determinants of local and regional differences in 8-oxoG-induced mutability are currently unknown. To uncover factors influencing the topology of 8-oxoG-induced mutations, we assessed spontaneous and KBrO_3_-induced 8-oxoG mutagenesis in human cell lines. KBrO_3_ exposure produced a SBS18-like substitution spectrum and a distinct never-before reported INDEL signature that we also observed in human cancers. KBrO_3_-induced 8-oxoG lesions occurred with similar sequence preference as KBrO_3_-induced substitutions, indicating that the reactivity of specific reactive oxygen species (ROS) dictates the trinucleotide motif specificity for 8-oxoG-induced mutagenesis. While 8-oxoG lesions occurred relatively uniformly across chromatin states and nucleosomes, 8-oxoG-induced mutations occurred more frequently in more compact regions of the genome, within nucleosomal DNA, and at inward facing guanines within strongly positioned nucleosomes. Cryo-EM structures of OGG1 bound to nucleosomes indicate that these effects originate from OGG1’s ability to flip outward positioned 8-oxoG lesions into the catalytic pocket with only minor alterations to nucleosome structure, while inward facing lesions occluded by the histone octamer are unrecognized. Mutation spectra from cells with DNA repair deficiencies revealed a hierarchical DNA repair network limiting 8-oxoG mutagenesis in human cells, where OGG1– and MUTY-mediated BER is supplemented by replication-associated factors participating in tolerance of 8-oxoG or derived repair intermediates (i.e. Pol η and HMCES). Surprisingly, analysis of transcriptional asymmetry of KBrO_3_-induced mutations demonstrated transcription-coupled repair of 8-oxoG in Pol η-deficient cells. Thus, radical chemistry, chromatin structures, and DNA repair processes combine to dictate the oxidative mutational landscape in human genomes.

## Introduction

Reactive oxygen species (ROS) react with nucleotide bases in DNA to form a variety of mutagenic lesions, including 8-oxoG adducts^1^. ROS are generated in cells by endogenous processes like lipid peroxidation and cell metabolism^2,3^ or through exposure to exogenous chemicals such as potassium bromate (KBrO_3_)^1,4^, a former food additive. During carcinogenesis, oncogene activation can also drive ROS formation through changes in metabolic oxidation^5^. Due to the prevalence of exogenous and endogenous agents that induce oxidative stress, oxidative lesions are the second most common DNA lesion following abasic sites (AP-sites)^6^. Moreover, mutations caused by oxidative damage are a common feature of human cancer genomes with single base substitution signature 18 (SBS18) hypothesized to arise from unrepaired 8-oxoG lesions. This signature occurs in ∼50% of sequenced human tumors and contributes an average of 300 mutations per genome^7^. SBS18 consists of G to T (and complementary C to A) substitutions and are distributed non-uniformly across the human genome^7,8^. What factors dictate the sequence and topological specificity of 8-oxoG-induced mutation in the human genome are unknown.

Mutations caused by oxidative damage are primarily prevented by the activity of base excision repair (BER), which eliminates 8-oxoG in duplex DNA. BER can be initiated by OGG1, a glycosylase that recognizes 8-oxoG across from cytidine (C) and excises the adducted base leaving an AP-site that is subsequently processed by downstream enzymes in the BER pathway^1,9^. If OGG1 fails to remove an 8-oxoG adduct, the adducted guanidine can mutagenicaly Hoogsteen base pair with an adenine inserted by multiple polymerases (i.e. Pol δ, η, κ, and ζ) during DNA synthesis^10–13^. These 8-oxoG:dA mispairs are identified by a second DNA glycosylase, MUTYH, which cleaves the adenine leaving an AP-site that is further processed by the BER pathway^14^. Due to their direct role in removing 8-oxoG or Hoogsteen paired adenines, loss of OGG1 and MUTYH results in increased mutation rates and an altered mutation spectrum consisting of higher numbers of G to T (and complementary C to A) substitutions^15–17^. This elevated mutation rate is believed to cause MUTYH-associated polyposis syndrome^16^, where individuals inheriting germline MUTYH mutations develop higher incidences of gastro-intestinal cancers throughout life^18–21^. Accordingly, human cells or cancer genomes with bi-allelic OGG1 or MUTYH mutations display respective SBS18^16,17,22^ and SBS36^23^ mutation signatures, both of which are dominated by C to A substitutions. In other organisms, like yeast, BER can be supported by additional mechanisms to limit 8-oxoG mutagenesis. For example, some eukaryotic trans-lesion synthesis polymerases, like Pol η, preferentially insert C across from 8-oxoG in template DNA^10,24,25^, allowing error-free bypass of the lesion^26^. In addition, yeast also utilize mismatch repair to remove adenines mispaired with 8-oxoG lesions^26–28^. Whether similar alternative 8-oxoG repair mechanisms limit mutagenesis in human cells remains to be determined.

Here, we unravel the sequence and topological determinants of 8-oxoG formation, repair, and mutagenesis, and decipher how different DNA repair and tolerance pathways coordinate to produce the oxidation-induced mutational landscapes observed in human cancer genomes. We found that KBrO_3_-treatment produces 8-oxoG lesions and mutations with similar, trinucleotide preferences indicating that ROS chemistry is the primary cause of mutational sequence specificity. Additionally, KBrO_3_ produced a unique INDEL signature that was also observed in human cancers, providing evidence that ROS induces other mutation types beyond the canonical G>T substitutions associated with 8-oxoG. Beyond sequence determinants, we identified that chromatin structure is a key topological determinant of oxidation-induced mutations in human cancer genomes. KBrO_3_-treatment resulted in mutations that were enriched in heterochromatin, nucleosome bound DNA, and bases facing the histone octamer, a similar phenomenon observed in the SBS18 mutational signature. Cryo-EM structures of OGG1 bound to nucleosomes containing 8-oxoG revealed the enzyme uses a DNA sculpting and base flipping mechanism for repairing 8-oxoG in the nucleosome, providing a mechanistic basis for the elevated mutational density at bases facing the histone octamer. Finally, analysis of mutation spectra from OGG1-, MUTYH-, Pol η-, and HMCES-deficient cells determined the human 8-oxoG repair network is hierarchically organized with OGG1 and MUTYH performing primary mutation avoidance while Pol η and HMCES functioning in secondary roles mediating tolerance of unrepaired 8-oxoG or 8-oxoG-derived AP sites. Subsequent analysis of Pol η-deficient cells has unveiled the presence of transcription-coupled repair of 8-oxoG on the transcribed strand of genes.

## Results

To characterize processes that modulate 8-oxoG mutagenesis and thereby dictate its distribution in human cells, we propagated independent clonal isolates of wild-type immortalized human retinal epithelial cells (RPE-1 hTERT) in the absence and presence of 250 µM KBrO_3_ for 100 days, to mimic exogenous exposure. Surviving clonal isolates were obtained following this outgrowth and genomic DNA was isolated for Illumina whole genome sequencing. Whole genome sequencing of outgrowth clones was compared to that of corresponding pre-outgrowth populations to identify mutations accumulated during propagation (Figure 1A). A total of 19684 and 128366 mutations were identified from untreated and KBrO_3_-treated cells, respectively, using the consensus calls of three probabilistic variant callers: VarScan2^29^, SomaticSniper^30^, and Strelka2^31^. Cells treated with KBrO_3_ had a 23.5-fold increase in substitutions and 3.35-fold increase in small insertion/deletion (INDEL) mutations per sequenced genome when compared to the non-treated cells (Figure 1B).

**Figure 1:**
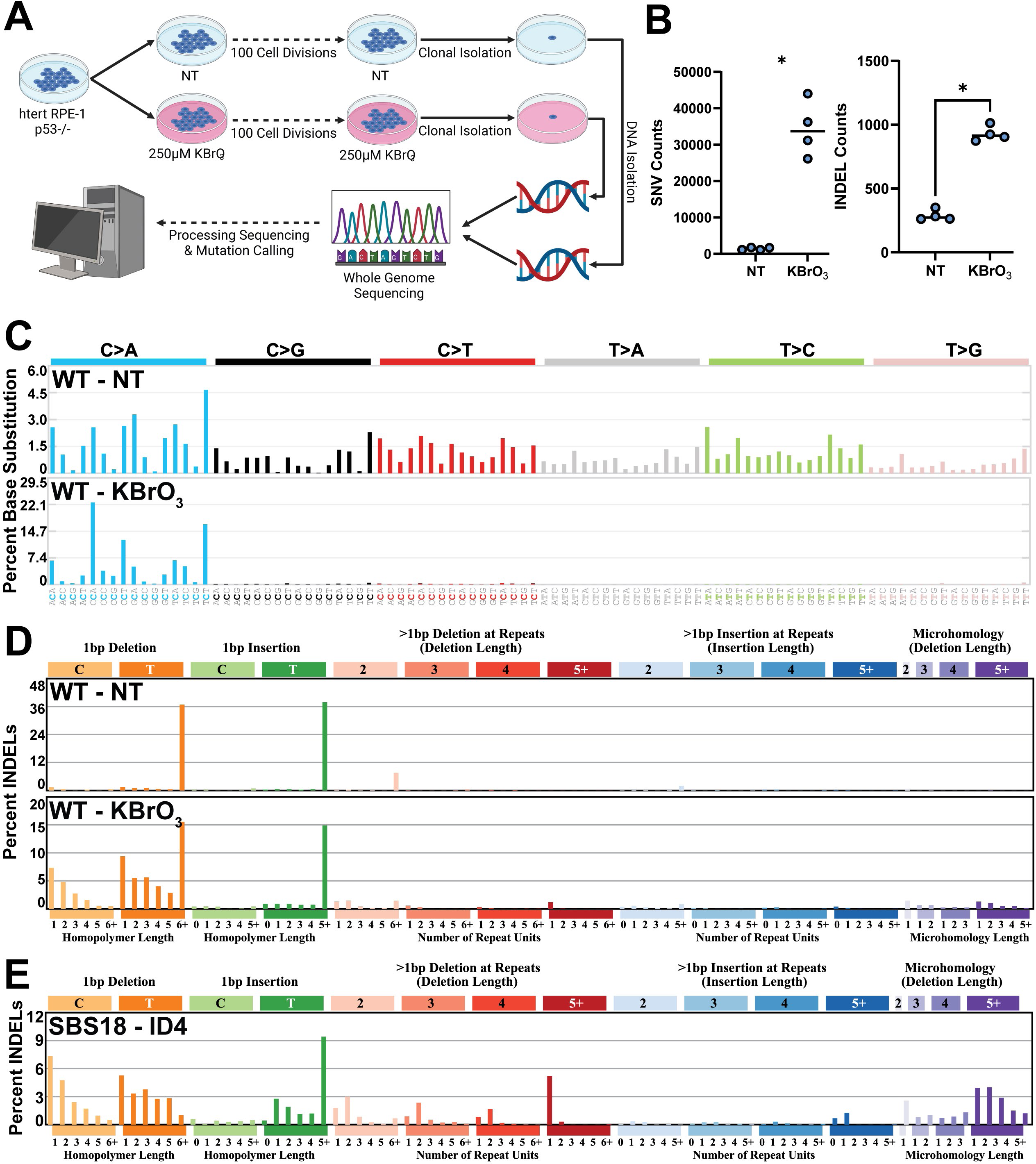
Mutagenesis in hTERT RPE-1 p53^−/−^ cells untreated and treated with 250µM KBrO_3_ after 100 cell divisions. **A**, Schematic of experimental conditions for mutation accumulation, clonal isolation, WGS, and mutation calling. **B,** Number of SNVs and INDELs per genome in untreated (NT) and KBrO_3_-treated (KBrO_3_) cells. Circles indicate independent genomes sequenced. Horizontal bars are median values. (* indicates p-value < 0.01 by Mann-Whitney U test) **C,** SNV and **D,** INDEL mutation signatures from treated and untreated genomes. **E,** De novo generated INDEL signature found in human cancers containing greater than 25% of total mutations attributed to SBS18. Total mutations involved in each signature are listed.

### Spontaneous and KBrO_3_-induced mutation spectra in human retinal pigment epithelial cells

Based upon this large increase in mutation load, we assumed most mutations within the KBrO_3_-treated cells were induced by ROS. We therefore produced a de novo KBrO_3_ specific mutation signature using SigProfilerExtractor^32^ set to detect 2 signatures from our dataset: one corresponding to the KBrO_3_-induced mutations (SBS96A) and the second representing the spontaneously acquired mutations during untreated outgrowth (SBS96B) (Supplemental Figure 1A,B). Deconvolution of these signatures into known COSMIC SBS signatures revealed that untreated RPE-1 cells contained a broad spectrum of base substitution mutations most consistent with SBS40, SBS5, and a small percentage of SBS18 (Supplemental Figure 1C), which are all consistent with spontaneously accumulated mutations during cell culture. Contrastingly, KBrO_3_-treated cells were dominated by C>A substitutions, as expected for mutations derived from unrepaired 8-oxoG (Figure 1C and Supplemental Figure 1D).

SigProfilerExtractor also identified 2 signatures for INDEL mutations. Untreated cells contained primarily 1 bp T insertions and deletions in long homopolymer runs and thus appears to be a combination of COSMIC INDEL (ID) signatures ID1 and ID2 (Figure 1D). These mutation signatures are associated with replication slippage events that would be expected to arise spontaneously through cell division^7^. KBrO_3_-treated cells also contained the ID1– and ID2-like mutations, though we observed a significant number of 1 bp deletions of C and T nucleotides (Figure 1D). These deletions were most common when not in homopolymer runs suggesting that they are likely induced by DNA damage and independent of polymerase slippage. While a 1 bp deletion of C bases could logically stem from error-prone replication past a KBrO_3_-induced 8-oxoG, the presence of a similar number of 1 bp T base deletions was surprising. KBrO_3_ largely produces 8-oxoG lesions through a reaction with glutathione that generates an unknown ROS^4,33^, and no T based lesion has been specifically identified. The sequences flanking 1 bp T deletions displayed a random distribution of C:G and A:T base pairs, indicating that the T deletions were unlikely to be collateral mutations caused by extended synthesis via deletion-prone TLS polymerases recruited to bypass an 8-oxoG^34^ (Supplemental Figure 2A).

We next evaluated whether a similar INDEL signature is potentially associated with endogenous ROS during cancer development. We obtained mutation calls for whole genome sequenced primary tumors from the International Cancer Genome Consortium (ICGC) and identified 68 tumors containing greater than 25% of their substitutions stemming from SBS18, meaning that ROS is a major mutagen in these samples. Subsequently, we utilized SigProfilerExtractor on these samples to produce de novo INDEL signatures. This analysis determined that the fourth-most abundant INDEL signature (constituting ∼12% of INDEL mutations) had high similarity to our KBrO_3_-induced INDEL signature (Figure 1E and Supplemental Figure 3; cosine similarity = 0.751). We also utilized mutationalpatterns.R^35^ to re-assign mutations in ICGC to the entire catalog of COSMIC signatures with the addition of our KBrO_3_-induced INDEL signature. Following this process, the number of mutations in our KBrO_3_-induced INDEL signature correlated with the number of SBS18 mutations (Supplemental Figure 2B), indicating that the two signatures are likely linked, and that endogenous ROS produces insertion/deletion mutations in addition to well characterized substitutions it tumors.

### Endogenous and exogenous ROS produce different mutation signatures

We next compared the KBrO_3_-induced SBS signature to SBS18, which is proposed to originate from ROS producing 8-oxoG in human cancers (Figure 2A). While both the KBrO_3_ and SBS18 signatures were dominated by C>A substitutions, the dominant sequence contexts at which mutations occur were different, producing a cosine similarity of only 0.812. This difference was most pronounced at the sequences CCA, CCT, GCA, and GCT. KBrO_3_-treatment produced a greater proportion of mutations at CCA and CCT sequences and a corresponding lower proportion of mutations in GCA and GCT sequences compared to SBS18. We wondered whether this difference in sequence specificity resulted from differences in the location of 8-oxoG formation caused by endogenous ROS compared to KBrO_3_-induced ROS. We therefore obtained CLAPS-seq reads, which identify the location of 8-oxoG lesions at single nucleotide resolution, from HeLa cells grown in the presence and absence of KBrO_3_^36^. Like mutations, the distribution of sequence contexts containing 8-oxoG lesions differed following KBrO_3_ treatment compared to untreated cells that formed 8-oxoG from endogenous processes (cosine similarity = 0.909) (Figure 2B). KBrO_3_-induced lesions occurred in contexts highly similar to KBrO_3_-induced mutations, except for a higher proportion of lesions occurring in the context of CCC, suggesting that these lesions may be either more accurately bypassed or preferentially repaired prior to mutagenesis. Endogenously generated 8-oxoG is also over-represented in the CCC context compared to mutations in SBS18. However, these lesions also occurred in CCT and GCC contexts more frequently than SBS18 mutations. Still, the difference of the KBrO_3_ and endogenous 8-oxoG proportional sequence contexts displayed a striking similarity to that of the KBrO_3_-induced mutations and SBS18, suggesting that differences in lesion formation largely account for the differences between the KBrO_3_ and SBS18 mutation signatures.

**Figure 2:**
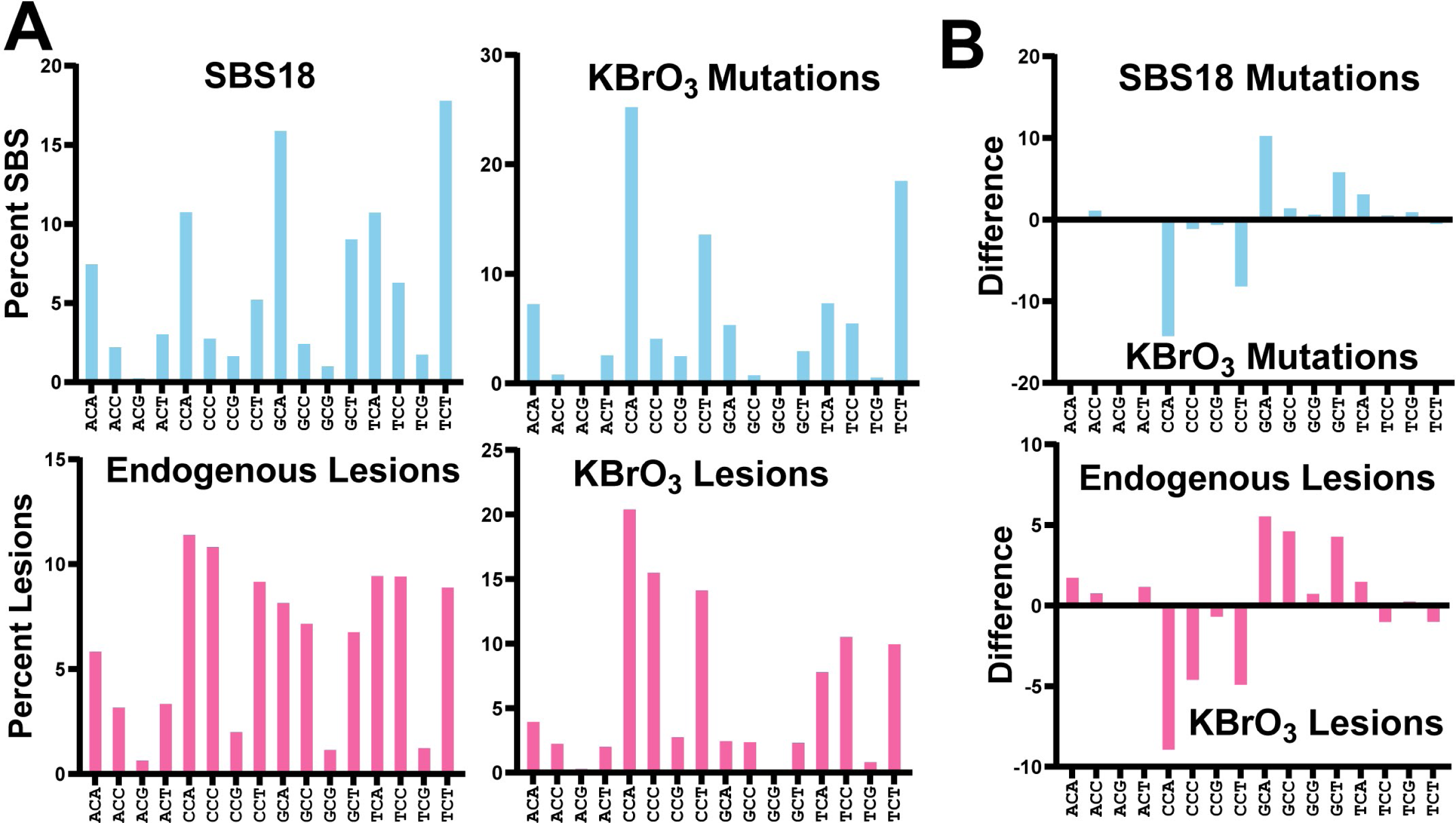
C>A mutation spectra and 8-oxodG lesion spectra in human cells under endogenous or KBrO_3_-induced DNA damage. **A**, Comparison of percentages of C>A trinucleotide mutation contexts from COSMIC SBS18 and KBrO_3_-induced mutations gave a cosine similarity of 0.812 and comparison with endogenous 8-oxodG lesion mapping gave a cosine similarity of 0.867. Comparison of KBrO_3_-induced mutations with KBrO_3_-induced 8-oxodG lesions gave a cosine similarity of 0.896. **B,** Differential bar graphs display discrepancies in trinucleotide contexts percentages between SBS18 and KBrO_3_ mutations has a cosine similarity of 0.908 compared to the differential bar graphs displaying discrepancies between trinucleotide contexts from endogenous 8-oxodG lesions compared to endogenous KBrO_3_-induced 8-oxodG lesions.

### BER reduces 8-oxoG mutations in solvent exposed, less chromatin compacted DNA

In human cancers, mutation densities caused by a variety of DNA damages are dictated in part by chromatin structure with heterochromatic regions having higher mutation rates arising from reduced DNA repair efficiency^37,38^. We sought to determine whether BER of 8-oxoG lesions was similarly impacted by chromatin resulting in a non-random distribution of mutations. We profiled 8-oxoG lesions and mutations in WT RPE-1 cells across chromatin states derived by Hidden Markov Modeling (HMM) of eight histone modifications and CTCF^39–41^ (Figure 3A). This modeling results in 15 chromatin states with different extents of euchromatic character. 3 of these states are associated with highly repetitive sequences and therefore were excluded from our analysis. By stratifying heterochromatin, promoters, enhancers, and transcribed regions into different states, we found that KBrO_3_-induced mutations decreased in less compact regions. Interestingly, 8-oxoG lesions were evenly distributed across all states indicating that inhibited DNA repair in heterochromatin likely underlies the higher mutation rates in these regions. As the repressive nature of heterochromatin is largely generated by tightly packed nucleosomes within these regions, we also profiled 8-oxoG mutations and lesions around strongly positioned nucleosomes within the human genome (Figure 3B). Consistent with our prior analysis, we observed 8-oxoGs formed relatively evenly across nucleosome bound regions. However, KBrO_3_-induced mutations oscillated with a ∼192 bp periodicity peaking within histone bound DNA, while lower mutagenicity was observed in linker DNA between nucleosomes. This finding is consistent with DNA repair being inhibited by tightly bound histones that obscure access to 8-oxoG during repair. Repair inhibition also extended within individual nucleosomes as KBrO_3_-induced mutations displayed a strong 10.3 bp oscillation (Figure 3B). The peaks of this oscillation occurred at inward facing nucleotides closest to the histones, whereas the troughs occurred at the most solvent exposed nucleotides. This result indicates that either 8-oxoG preferentially forms at histone proximal nucleotides or that repair of 8-oxoG by BER is more efficient at solvent exposed lesions. 8-oxoG lesions displayed a similar pattern as KBrO_3_-induced mutations, but with a significantly lesser amplitude. We therefore conclude that efficient BER at outward facing bases is the primary factor influencing 8-oxoG mutations within nucleosomes, though a subtle lesion formation preference may exist. To determine if other DNA proteins beyond histones can block the repair of 8-oxoG, we profiled KBrO_3_ mutations and lesions at active transcription factor binding sites (Figure 3C). We found that neither mutations nor lesions were elevated at these sites in contrast to other types of DNA damage like CPDs^42^, suggesting that the inhibition of 8-oxoG repair is specific for nucleosome structure.

**Figure 3:**
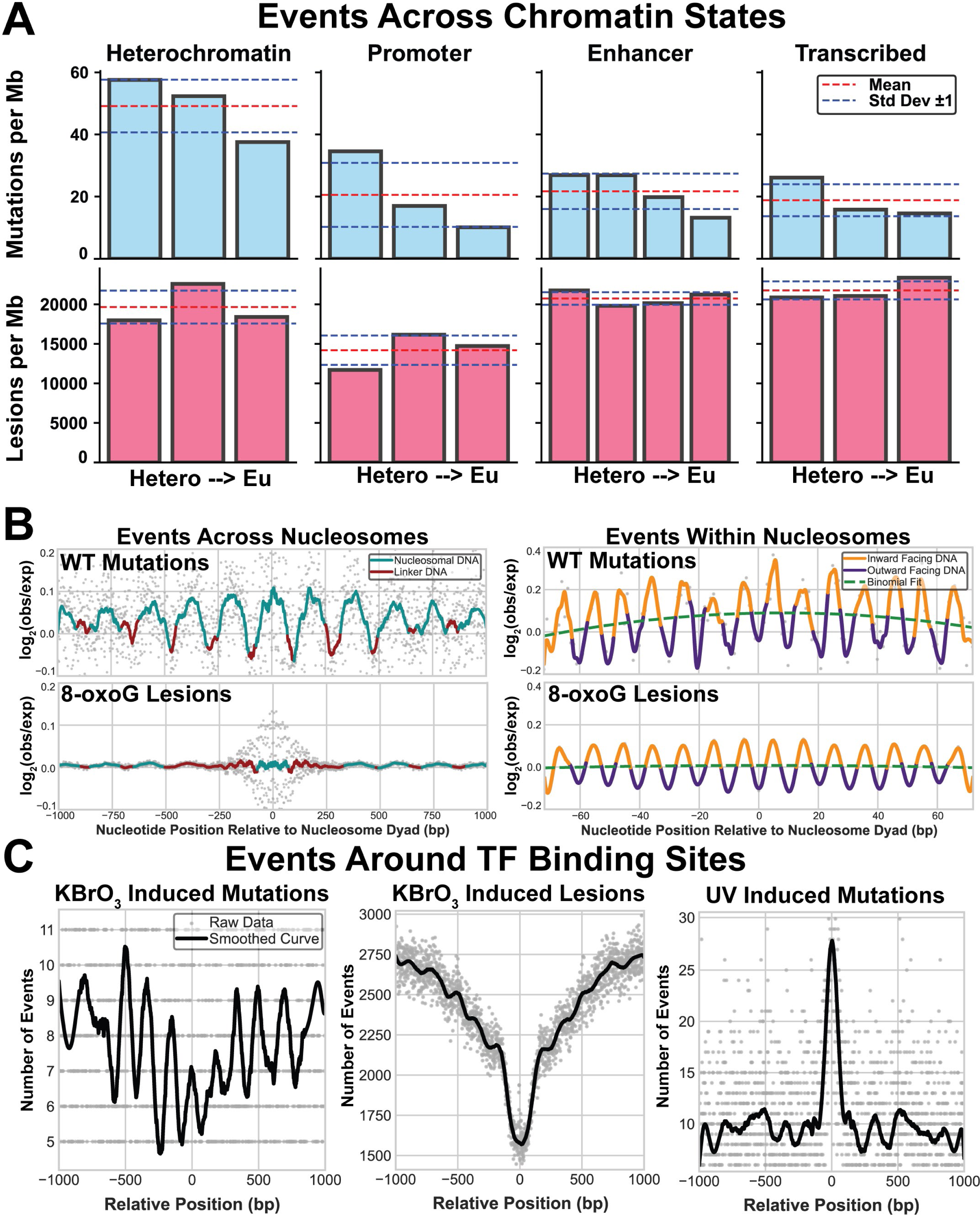
Chromatin state, nucleosome binding, and transcription factor binding’s impact on 8-oxodG mutagenesis and lesion formation. **A**, Four binned broad regions are dictated by the ChromHMM map and within each group are sorted from left to right as being more heterochromatic to more euchromatic. Mutations are represented in the top set of bars in blue and lesions are represented in the bottom set of bars in pink. **B,** The left graphs represent translational periodicity of log2(observed/expected) of events between nucleosomes with mutations on top and lesion on the bottom. Nucleosome bound DNA is represented in blue and linker DNA is represented in red. The right two graphs represent the rotational periodicity of the log2(observed/expected) of events within the nucleosome where DNA that is inward facing relative to the nucleosome is displayed in gold while outward facing relative to the nucleosome is displayed in purple. A binomial fit of the data is overlayed in a dashed green line. In both figures, actual data points are displayed in gray. **C,** Number of events are plotted relative to the TF binding midpoint for mutations on the left and lesions on the right. Original data points are displayed in gray and a smoothed curve is overlayed in black.

The mapping of 8-oxoG lesions and KBrO_3_-induced mutagenesis indicate that 8-oxoG undergoes preferential repair at solvent-exposed positions in the nucleosome (Figure 3B). Consistent with these findings, previous work identified that the DNA glycosylase OGG1 excises 8-oxoG from solvent-exposed positions more efficiently than histone-occluded positions in recombinant nucleosomes *in vitro* ^43,44^. To obtain mechanistic insight into the preferential repair of solvent-exposed 8-oxoG in the nucleosome, we determined a 3.3 Å cryo-EM structure of OGG1 bound to a nucleosome containing a solvent-exposed 8-oxoG at superhelical location (SHL)_−6_, referred to as OGG1-8oxoG-nucleosome core particle (NCP)_−6_ (Figure 4A, Supplemental Figures 4-6, and Supplemental Table 1). We utilized a catalytically dead variant of OGG1 (K249Q) that maintains the ability to specifically recognize 8-oxoG ensuring we captured an 8-oxoG substrate recognition complex^45,46^. The local resolution of the nucleosome was 3 – 4 Å and the local resolution of OGG1 was 5 – 7 Å (Supplemental Figure 5F), which was sufficient to unequivocally dock a previously determined high-resolution X-ray crystal structure of OGG1 (PDB: 1EBM)^45^ into the cryo-EM map (Supplemental Figure 5H).

**Figure 4:**
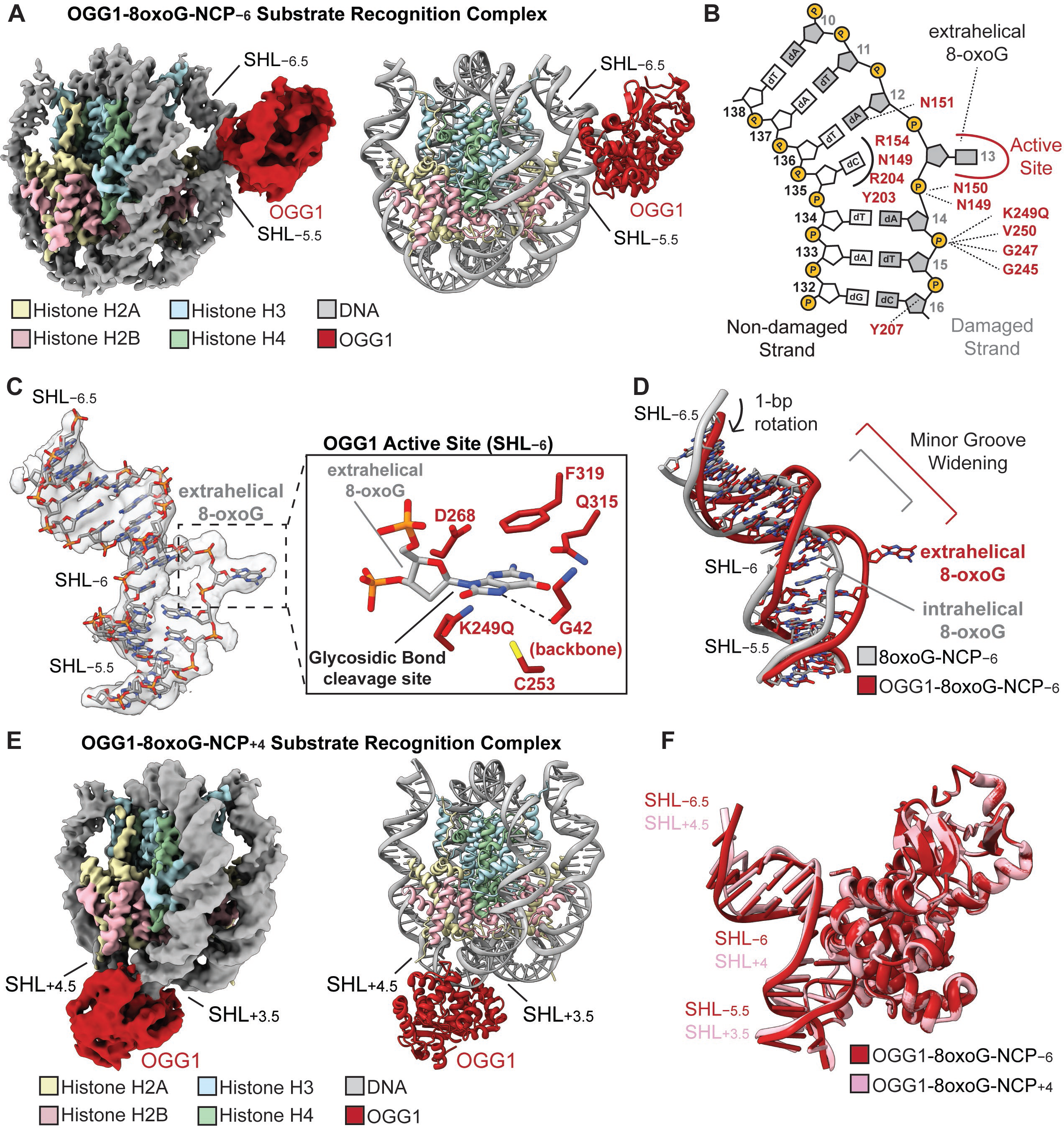
Single particle analysis of OGG1-8oxoG-NCP_−6_. **A**, The 3.3 Å OGG1-8oxoG-NCP_−6_ composite cryo-EM map (left) and cartoon representation of the OGG1-8oxoG-NCP_−6_ model (right). **B,** A diagram representing the interactions between OGG1 and the nucleosomal DNA in the OGG1-8oxoG-NCP_−6_ complex identified using PLIP^85^. **C,** Focused view of the nucleosomal DNA at SHL_−5.5_ to SHL_−6.5_ showing the extrahelical 8-oxoG at SHL_−6_. The segmented density for the nucleosomal DNA in the OGG1-8oxoG-NCP_−6_ composite cryo-EM map is shown in transparent grey. An inset of the OGG1 active site is shown, highlighting key amino acids important for 8-oxoG recognition and excision. **D,** Structural comparison of the nucleosomal DNA (SHL_−5.5_ to SHL_−6.5_) in the OGG1-8oxoG-NCP_−6_ complex and 8oxoG-NCP_−6_, highlighting the structural changes in the nucleosomal DNA induced by OGG1 binding. **E,** The 3.6 Å OGG1-8oxoG-NCP_+4_ composite cryo-EM map (left) and cartoon representation of the OGG1-8oxoG-NCP_+4_ model (right). **F,** Structural comparison of OGG1 and the nucleosomal DNA (SHL_−5.5_ to SHL_−6.5_) in the OGG1-8oxoG-NCP_−6_ and OGG1-8oxoG-NCP_+4_ complexes, highlighting the similarities in 8-oxoG recognition at both positions.

In the OGG1-8oxoG-NCP_−6_ substrate recognition complex, OGG1 is engaged with ∼5 base pairs of nucleosomal DNA at SHL_−6_, which buries ∼1086 Å^2^ of surface area (Figure 4A). The interaction of OGG1 with the nucleosomal DNA is mediated by a network of non-specific interactions with the phosphate backbone of the damaged nucleosomal DNA strand, as well as extensive contacts with the orphan cytosine and 8-oxoG (Figure 4B and 4C). Interestingly, we did not observe a direct interaction between OGG1 and the histone octamer, indicating that nucleosome binding by OGG1 is primarily driven by the interactions with nucleosomal DNA. At the center of the OGG1 binding footprint lies the nucleosomal 8-oxoG, which has been evicted from the DNA helix and positioned into the OGG1 active site (Figure 4B and 4C). In this conformation, the extrahelical nucleosomal 8-oxoG is positioned in proximity to key OGG1 amino acid residues that are important for 8-oxoG binding specificity (G42 carbonyl), stabilization of the extrahelical 8-oxoG (C253, F319, and Q315) and 8-oxoG excision (K249Q and D268) (Figure 4C)^45^. Cumulatively, this data shows OGG1 is in a conformation poised for 8-oxoG excision.

To position the 8-oxoG into the catalytic active site, OGG1 binding induces significant structural changes in the nucleosomal DNA during 8-oxoG recognition. These structural changes include a 1 bp rotation of the nucleosomal DNA, significant minor groove widening at SHL_−6_, and nucleosomal DNA bending around SHL_−5.5_ to SHL_−6.5_ when compared to 8oxoG-NCP_−6_ (Figure 4D). The OGG1-induced minor groove widening and nucleosomal DNA bending facilitate extrusion of the 8-oxoG from the nucleosomal DNA into the OGG1 active site (Figure 4C). Ultimately, the mode of 8-oxoG recognition and the OGG1-induced structural changes in the nucleosomal DNA are similar to those seen for OGG1 bound to 8-oxoG in non-nucleosomal DNA (RMSD_DNA_ – 1.621) (Supplemental Figure 7A)^45^, indicating a conserved 8-oxoG recognition mechanism in chromatin and non-chromatinized DNA.

To determine whether OGG1 uses the same mechanism for 8-oxoG recognition at different translational locations in the nucleosome, we determined a 3.6 Å cryo-EM structure of OGG1 K249Q bound to a nucleosome containing a solvent-exposed 8-oxoG at SHL_+4_, referred to as OGG1-8oxoG-NCP_+4_ (Figure 4E, Supplemental Figures 8-10, and Supplemental Table 1). The general mechanism of nucleosome binding and 8-oxoG recognition by OGG1 at SHL_+4_ are similar to those observed for OGG1 bound to 8oxoG at SHL_-6_ (Supplemental Figure 7B,C). However, OGG1 binding induces modest structural rearrangements in the nucleosomal DNA during 8-oxoG recognition at SHL_+4_, which includes minor groove widening of the nucleosomal DNA without significant DNA bending. The lack of OGG1-induced DNA bending is likely due to the inherently bent conformation of the nucleosomal DNA near SHL_+4_ (Supplemental Figure 7D). Despite these subtle differences, the final conformation of OGG1 and the nucleosomal DNA in the OGG1-8oxoG-NCP_+4_ and OGG1-8oxoG-NCP_−6_ structures are extremely similar (Figure 4E and Supplemental Figure 7A). This data strongly suggests that OGG1 uses the same general mechanism for 8-oxoG recognition and repair at solvent-exposed positions throughout the nucleosome. Notably, the structural changes observed during the recognition of solvent-exposed 8-oxoG by OGG1 are incompatible with binding 8-oxoG proximal to the histone octamer, as this would result in significant clashes between OGG1 and the core histone octamer (Supplemental Figure 7E,F). Together, this data provides a strong structural rationale for the preferential repair of solvent-exposed 8-oxoG in the nucleosome *in vitro* and *in vivo,* and the elevated levels of KBrO_3_-induced mutagenesis at nucleotides proximal to the histone octamer (Figure 3B)^43,44^.

### Replication and transcription-associated mechanisms limit 8-oxoG mutagenesis

In various species, 8-oxoG mutagenicity is limited by multiple, redundant DNA repair and damage tolerance pathways. This includes the activities of OGG1-initiated BER, MutY-initiated BER, mismatch repair, nucleotide excision repair (NER), and accurate TLS bypass by DNA polymerase η. We therefore compared spontaneous and KBrO_3_-induced mutation spectra among WT human cell lines and those lacking OGG1^17^, MUTYH^17^, Pol η^47^, or HMCES, a recently identified replication-associated factor that participates in bypass of ssDNA lesions^48–52^ and protects cells from cytotoxicity associated with KBrO_3_ exposure^48,51^. Loss of OGG1, MUTYH, and HMCES resulted in moderate ∼2 to 3-fold increases in the amount of spontaneously acquired mutations per genome compared to corresponding WT lines, while Pol η-deficiency failed to increase spontaneous mutagenesis (Supplemental Figure 11A). RPE-1 cells lacking HMCES maintained similar spectra suggesting the increased spontaneous mutation load results from a general reduction in error-free lesion bypass (Supplemental Figure 11B). Contrastingly, loss of either OGG1 or MUTYH produced spectra consisting primarily of SBS18-like mutations, indicating these glycosylases are the primary mechanism for preventing 8-oxoG mutagenesis and HMCES is likely involved more generally in lesion bypass (Supplemental Figure 11B).

To directly evaluate the role of Pol η and HMCES in 8-oxoG bypass, we compared mutation spectra from RPE-1 knockout lines following prolonged KBrO_3_ exposure to those in WT RPE-1 cells (evaluated in Figure 1). Deficiency in Pol η or HMCES resulted in 1.5– and 1.3– fold increases in total KBrO_3_-induced mutations, respectively (Figure 5A). These mild increases impacted both substitutions and INDEL mutation types (Figure 5A) but likely underestimates the true augmentation of KBrO_3_ mutagenesis as HMCES^−/−^ cells showed a significant growth delay upon initial KBrO_3_ exposure likely resulting in fewer cell divisions for these lines. The substitution spectra of Pol η– and HMCES-deficient cells were nearly identical to that of KBrO_3_-treated WT cells (Figure 5B), indicating that both enzymes likely participate in some form of error-free bypass of 8-oxoG or derived repair intermediates. Consistent with this mechanism, neither gene disruption changed the impact of chromatin compaction on 8-oxoG induced substitution frequency (Supplemental Figures 12 and 13), suggesting they are primarily operating in contexts without nucleosome involvement. INDEL spectra from KBrO_3_-treated Pol η– and HMCES-deficient cells, were also like that of WT cells, except for several subtle differences. HMCES^−/−^ cells displayed a small increase in 1 bp T insertions at shorter homopolymer lengths, while Pol η-deficiency resulted in a general loss of 1 bp T insertions and a preference for 1 bp C or T deletions occurring in 2-3 bp homopolymer repeats (Figure 5C). Ultimately, these differences in mutation spectra, particularly for INDELs, further supports roles for both Pol η and HMCES in ROS-induced lesion bypass, although these mechanisms of limiting mutation appear to function as a secondary level of protection behind the activities of OGG1 and MUTYH.

**Figure 5:**
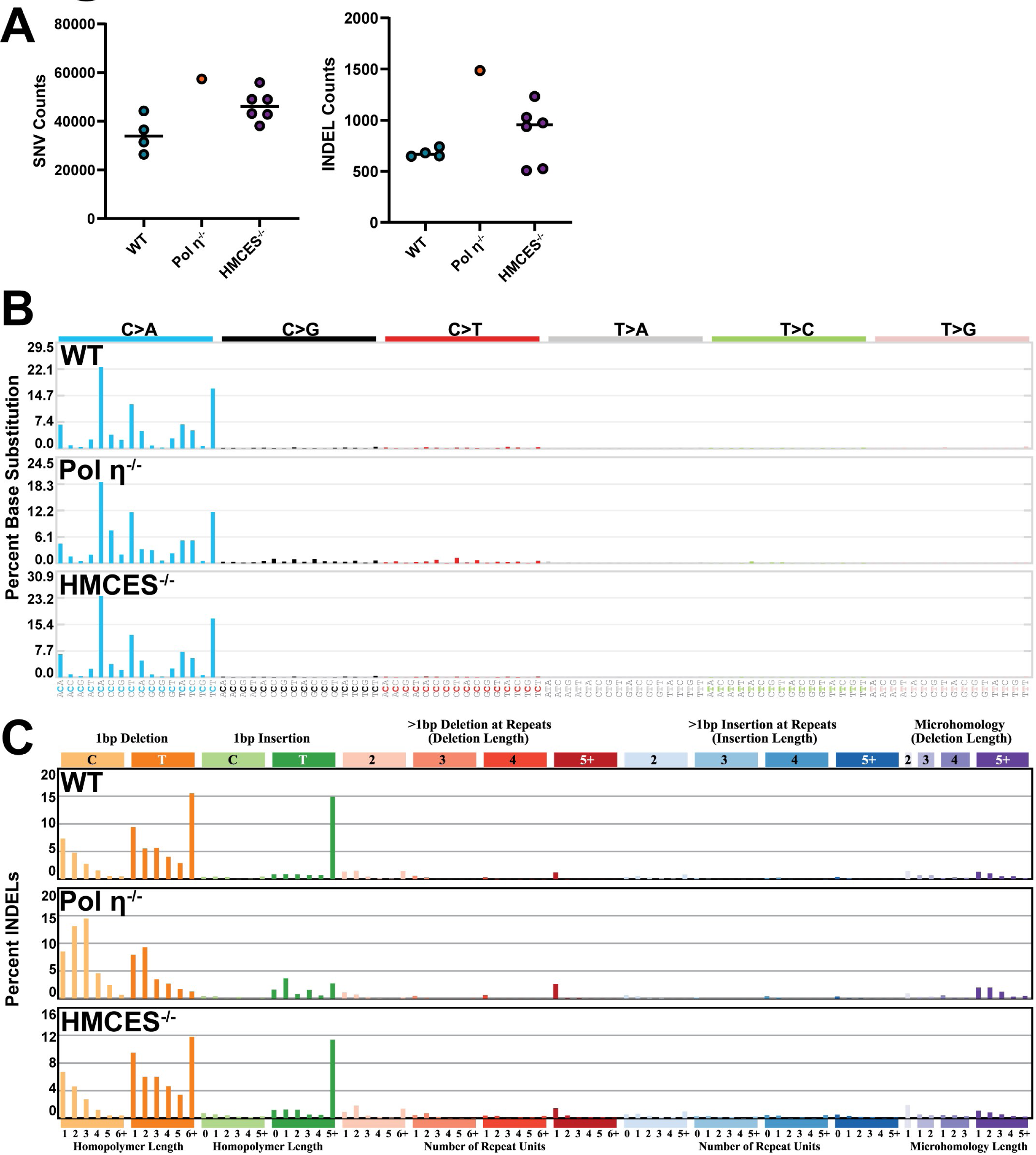
KBrO_3_-induced mutagenesis in human WT, POLH^−/−^, or HMCES^−/−^ RPE-1 cells lines. **A**, Number of SNVs and INDELs per genome in KBrO_3_-treated for WT, POLH^−/−^, and HMCES^−/−^ RPE-1 cells lines. Circles indicate independent genomes sequenced and horizontal bars are median values. **B,** SNV and **C,** INDEL mutation signatures from treated and untreated genomes. Total mutations involved in each signature are listed.

To determine whether mismatch repair (MMR) limits 8-oxoG mutagenesis in human cells, we evaluated replication strand asymmetry of KBrO_3_-induced mutations using AsymTools2 software^53^ that determines leading and lagging strand association of mutations based upon the directionality of the replication fork movement in the mutated region. Applying this analysis to CLAPS-seq reads in KBrO_3_-treated WT cells revealed an equal distribution of 8-oxoG lesions on the leading and lagging template strands (Figure 6A). 8-oxoG-induced mutations however displayed slightly more G>T substitutions on the leading strand suggesting that a DNA repair process (potentially MMR) preferentially removes 8-oxoG from the lagging strand template. HMCES deletion exacerbated this effect, indicating that HMCES may favor bypass of leading strand lesions. Interestingly, loss of Pol η removed the replication strand asymmetry. This indicates that Pol η likely mediates error-free bypass of 8-oxoG in the lagging strand template in human cells. Similar results indicate Pol η TLS functions preferentially during lagging strand synthesis for bypass of UV photoproducts in human melanomas and fibroblasts^54^, suggesting a general lagging strand preference for this TLS polymerase.

**Figure 6:**
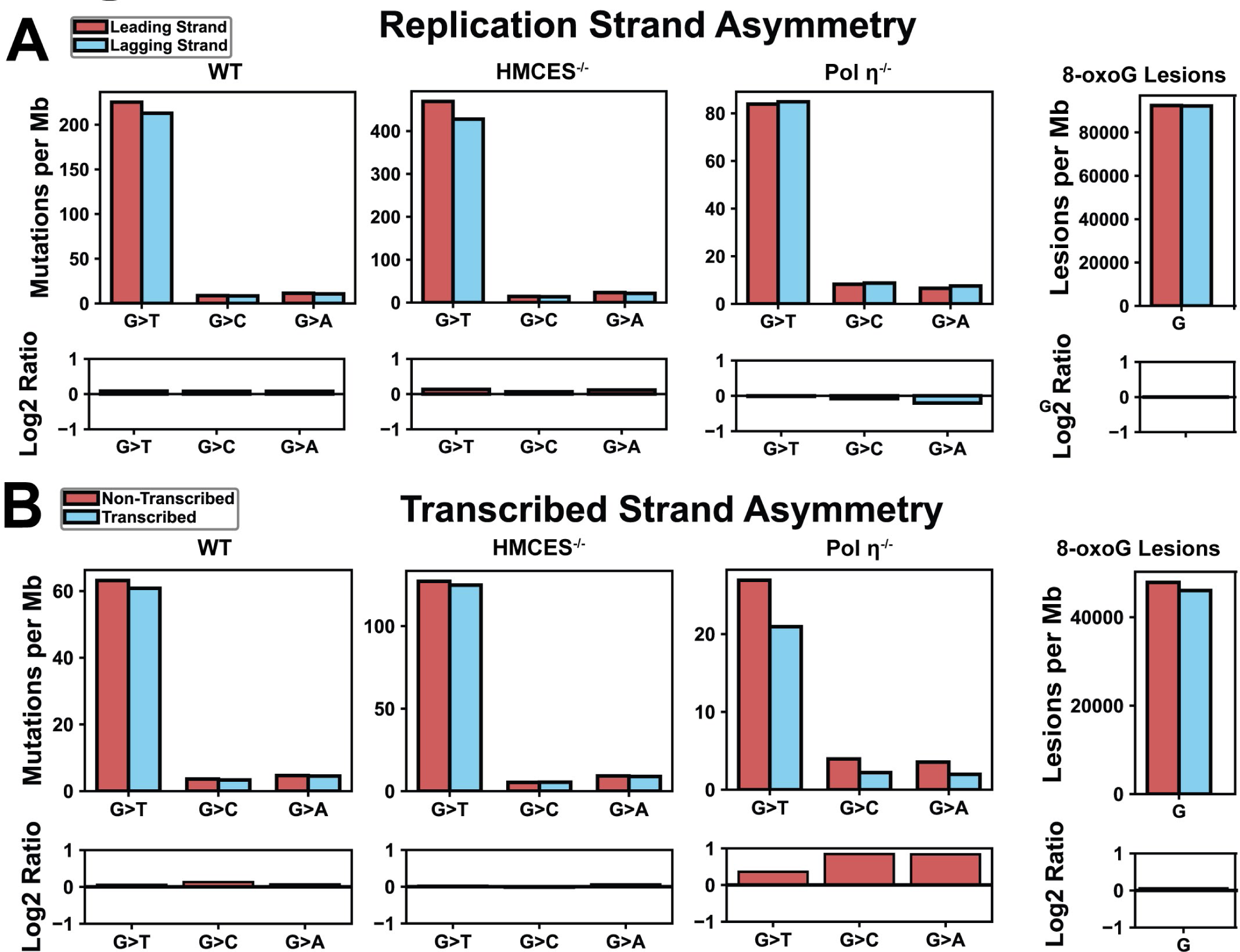
KBrO_3_-induced 8-oxodG mutation and lesion strand bias on leading/lagging and transcribed/non-transcribed strands in human cells. Red bars represent G>T mutations on the **A,** leading strand while blue bars represent G>T mutations on the lagging strand. **B,** Red bars represent G>T mutations on the non-transcribed strand and blue represents G>T mutations on the transcribed strand. The bar graph below each plot represents the log2(red/blue) value of the for the event per MB bars.

We also assessed whether KBrO_3_-induced mutations displayed transcriptional asymmetry, which would be indicative of transcription coupled repair of 8-oxoG. Transcription-coupled NER and BER have been suggested to be involved in 8-oxoG removal^55,56^. However, little transcriptional mutation asymmetry has been reported for ROS-associated SBS18 mutations^8^, suggesting that transcription coupled repair of 8-oxoG may be limited. We found that G>T substitutions in WT and HMCES^−/−^ cells only slightly favored the non-transcribed strand of genes (Figure 6B). This bias was also observed in CLAPS-seq reads for 8-oxoG lesions, suggesting this effect is caused by a preference in lesion formation instead of a repair process. Transcriptional asymmetry in Pol η-deficient cells, however, was very pronounced and impacted all G>T, G>C, and G>A substitution types. The effect size of this asymmetry is on par with other mutational processes limited by transcription-coupled nucleotide excision repair^53^ and provides strong evidence that transcription-coupled repair can remove 8-oxoG lesions. However, the impact of transcription-coupled repair in mutational data is obscured by the error-free bypass of the lesion by Pol η.

## Discussion

### Mutations from endogenous and exogenous ROS

We observed that long-term treatment of RPE-1 cells with the oxidant KBrO_3_ increases substitutions and produces a mutational signature like COSMIC SBS18 that is proposed to be caused by endogenous ROS. Both signatures are composed of C>A mutations, however, the preferred trinucleotide sequences these substitutions occur in differ. Specifically, KBrO_3_ exposure produced an over-representation of substitutions at CCA and CCT and under-representation at GCA, and GCT compared to SBS18. Other studies have demonstrated the same KBrO_3_ signature for both long term exposure in RPE-1 cells^47^ or short term exposure of human iPSCs^57^, indicating that differences in exposure protocol and/or cell lines are not responsible for the differences in SBS18 and KBrO_3_-induced substitution specificity. Additionally, CLAPs-seq mapping of KBrO_3_-induced 8-oxoG lesions in HeLa cells produced similar sequence preferences as KBrO_3_-induced mutations, providing evidence that the oxidizing agent can dictate the sequences most likely to form 8-oxoG. Interestingly, CCA and CCT mutation contexts correspond to trinucleotides with low vertical ionization potential (VIP), which sensitizes these motifs (i.e. TGG and AGG sequences) to long-range guanine oxidation by charge transfer^58^.

Reciprocally, TGC and AGC have higher VIPs, indicating that GCA and GCT sequences would have fewer mutations produced by this mechanism. This correlation suggests that KBrO_3_ may induce more guanine oxidation through charge transfer than endogenous ROS, leading to an KBrO_3_ specific 8-oxoG mutation pattern. An alternative possibility is that specific reactive oxygen species produce mutations at different sequences. While the reactive oxygen species that is generated by KBrO_3_ is unknown, KBrO_3_ requires the presence of a reducing agent, like glutathione, to create 8-oxoG and the lesion’s formation is insensitive to traditional cellular ROS scavengers, indicating KBrO_3_ produces 8-oxoG by a different mechanism than endogenous ROS^4^. By extension, the DNA damage induced by endogenous ROS could result from the combined activity of multiple different species (e.g. peroxide, superoxide, etc.), which may all have different sequence preferences in forming 8-oxoG. In the future, utilizing human cell systems to determine the mutation signatures of individual endogenous reactive oxygen species will be beneficial in determining which sources of ROS are most relevant for inducing mutation in human cancers.

KBrO_3_ exposure primarily produces DNA damage in the form of 8-oxoG^33^, which canonically produces G>T substitutions through Hoogsteen base pairing of 8-oxoG with dA^59^ during DNA synthesis. We were therefore surprised to observe that KBrO_3_ treatment also increased INDEL mutations and that a large percentage of these mutations occurred at A:T pairs. While a 1 bp deletion of C bases could logically stem from error-prone replication past an KBrO_3_-induced 8-oxoG, the presence of a similar number of 1 bp T base deletions suggested that these mutations were caused either through collateral mutagenesis^34^ adjacent to an 8-oxoG or by a second KBrO_3_-induced DNA lesion. T deletions lacked an enrichment of C:G base pairs flanking the mutation, indicating that they were unlikely incurred as collateral mutations during 8-oxoG bypass. The lack of a reasonable connection of T deletions to 8-oxoG suggests that the ROS created by KBrO_3_ also causes at least one other mutagenic DNA lesion and targets T or A bases, such as thymine glycol^4,60^. Our analysis of INDEL signatures in human tumors displaying high levels of SBS18 mutations indicated that a similar INDEL process occurs in these tumors. Moreover, the number of SBS18 mutations correlates with the number of mutations attributed to our novel INDEL signature, both suggesting that endogenous ROS produces these mutations. Additional research is needed to identify the specific DNA lesions causing this signature.

### Oxidation-induced mutagenesis within the context of chromatin

Beyond the sequence specificity of the ROS created by KBrO_3_, chromatin structure is a major influence on the density of 8-oxoG-induced mutations. Higher mutation densities were observed in more compact regions of the genome, which likely stems from the density of nucleosomes within these regions. Within nucleosomes, mutations had a 10.3 bp periodicity when treated with KBrO_3_, occurring primarily on nucleotides proximal to the histone octamer. This suggests that DNA repair mechanisms may be excluded from the inward facing positions of the nucleosomal DNA. This is in stark contrast to UV-induced cyclo-pyrimidine dimer positioning at outward facing nucleotides at nucleosomes due to preferential lesion formation at these sites^61^. Indeed, our structural data demonstrates that OGG1 accesses outward facing 8-oxoG in the nucleosome by sculpting nucleosomal DNA and flipping the base into its active site for catalysis, similar to what was previously observed for the DNA glycosylase AAG^62^ and APE1^63^. This leaves inward facing 8-oxoG in the nucleosome more prone to mutation, as OGG1 lacks the ability to recognize 8-oxoG in these positions without massive structural changes in nucleosome structure. Repair of these sites is likely significantly delayed and may require active nucleosome remodeling in response to DNA damage by additional cellular factors, such as the BER-associated nucleosome remodeler ALC1^64^. Interestingly, other protein-DNA interactions appear to have little to no impact on 8-oxoG-induced mutagenesis. Transcription factors bound to gene promoter regions produce no change in the density of KBrO_3_-induced mutations in contrast to other types of DNA damage, like cyclobutane pyrimidine dimers. 8-oxoG lesions at these sites are greatly reduced compared to neighboring DNA, suggesting that they are rapidly repaired. However, we are unable to exclude the possibility that transcription factor binding protects their binding sites from formation of 8-oxoG.

### An expanded 8-oxoG repair network in human cells

Our data indicates at least three layers of 8-oxoG repair activities influence the mutagenicity of KBrO_3_ exposure (Figure 7). OGG1 and MUTYH activities provide the “first line of defense” against 8-oxoG mutagenesis as their deletion results in spontaneous mutator phenotypes displaying SBS18-like mutation spectra. These BER activities are in turn backed up by the replication-associated damage tolerance mechanisms of Pol η and HMCES. Human Pol η appears to specifically participate in the bypass of 8-oxoG directly as its deletion produces no change in spontaneous mutagenesis, indicating low utilization against most forms of endogenous DNA damage. However, Pol η-deficient cells had increased mutation load in response to KBrO_3_-treatement, indicating it participates in the accurate bypass of 8-oxoG generally across the human genome. The 1.5-fold increase in mutation load in KBrO_3_-treated Pol η-deficient cells is significantly lower than the impact of Pol η loss in yeast^24^. This suggests either lesser reliance on Pol η due to redundancy with other human DNA polymerases in 8-oxoG bypass, or less accurate Pol η bypass in humans compared to yeast. Supporting the latter, biochemical experiments have shown human Pol correctly inserts dC across from 8-oxoG to a lesser extent than yeast Pol η^24^.

**Figure 7:**
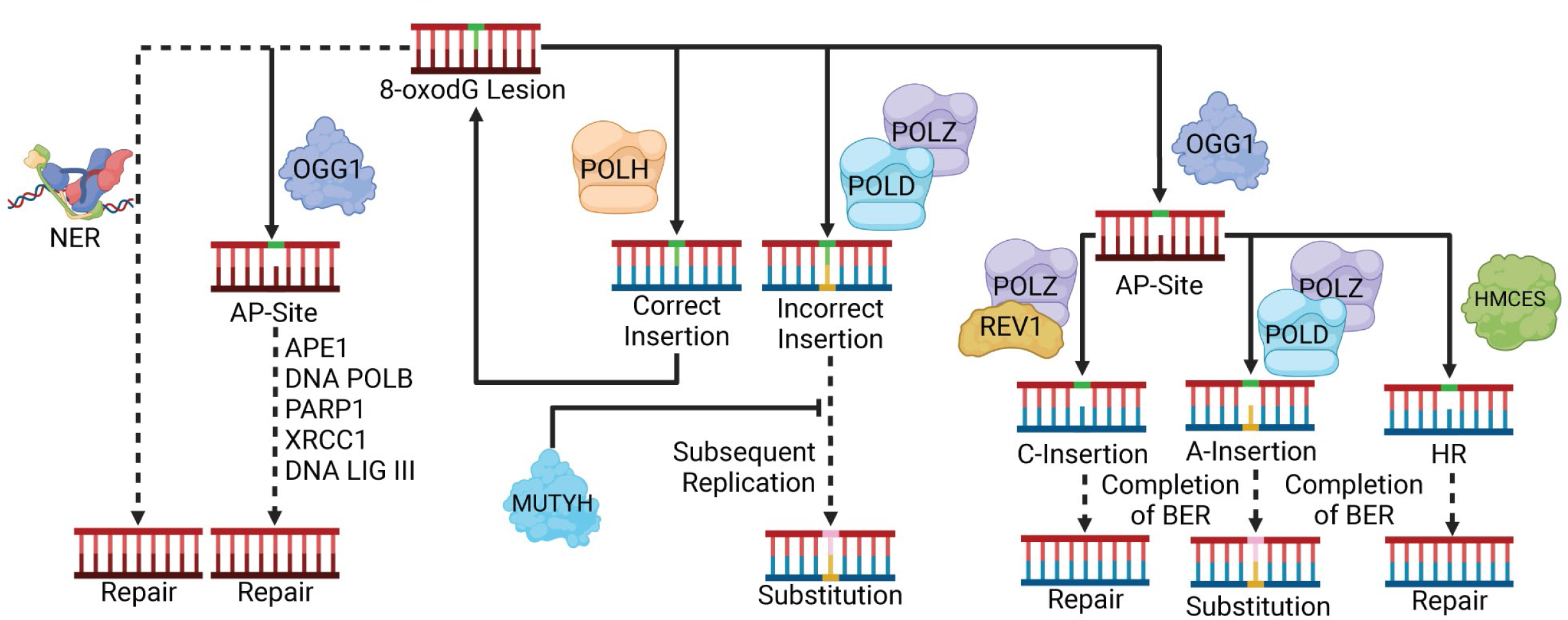
Mechanisms that limit KBrO_3_-induced 8-oxodG mutagenesis in human cells. Mutations from 8-oxoG can be limited in human cells by OGG1– and MUTYH-initiated BER. Secondary limits on mutagenesis include Pol η, HMCES, and transcription-coupled repair. Image created with Biorender.com under agreement number DX26OBAXYI.

HMCES-deficiency also resulted in elevated KBrO_3_-induced mutagenesis indicating it also protects against 8-oxoG mutagenesis, consistent with the previously reported sensitivity of HMCES^−/−^ cells to KBrO_3_^48^. This increase in mutagenesis in response to 8-oxoG was surprising considering HMCES was previously demonstrated to prevent mutagenesis^48,52,65,66^ by cross-linking to AP-sites^48^ and interacting with TLS polymerases^52^. Moreover, KBrO_3_-treated HMCES-deficient cells displayed a mutation spectrum identical to KBrO_3_-treated wildtype cells suggesting that HMCES is aiding in error-free bypass of 8-oxoG or derived repair intermediates. Such a mutation spectrum could be reconciled with HMCES’s known biochemical activity toward AP-sites if OGG1 removal of 8-oxoG can be uncoupled from the rest of BER, resulting in AP-sites occurring specifically at dG nucleotides. Subsequent TLS-based bypass of these AP-sites could either be error-free (in the case of REV1-mediated bypass via C-insertion) or produce C>A substitutions (by A-rule AP-sites site bypass), which would recapitulate the KBrO_3_ substitution signature. The error-prone bypass of AP-sites may also cause the oxidation-induced 1 bp deletions we identified and that are elevated in HMCES^−/−^ cells^67^. By analogy with Ung1 generated AP-sites sites from dU in the lagging strand template^68^, uncoupling of OGG1 glycosylase activity from BER would be expected to occur more frequently at sites of DNA replication (where HMCES functions) as the glycosylase could recognize 8-oxoG in ssDNA, but subsequent steps of BER would be inhibited do to the lack of a complementary DNA strand. HMCES loss did result in a modest increase in spontaneous mutagenesis with a broad mutation spectrum, which would be consistent with more error-prone AP-site bypass.

Mutational strand asymmetries in KBrO_3_-treated cells revealed a third level of protection from 8-oxoG-induced mutation. We observed minor differences in the mutational burden associated with the non-transcribed and transcribed strands of genes in both wildtype and HMCES deficient cells treated with KBrO_3_. In Pol η−deficient cells, however, there is a substantial decrease in the mutational burden on the transcribed strand. This may indicate that transcription-coupled repair processes are important for preventing mutagenesis from oxidative damage in the absence of Pol η. Alternatively, TC processes (TC-NER or TC-HR) may always be important or engaged for oxidative damage and the phenotype is exacerbated by the absence of Pol η. Previous studies indicate that CSB, a major component of TC-NER, can be recruited to transcription sites upon oxidative damage and is important for fitness following oxidative damage^56,69^. In contrast to transcriptional asymmetry in Pol η-deficient cells, only modest replication asymmetry toward the leading strand was observed for KBrO_3_-induced mutations in wild type cells, which could be indicative of preferential MMR repair of 8-oxoG on the lagging strand^70,71^. However, leading strand replication strand-bias was exacerbated in HMCES^−/−^ cells and removed in Pol η-deficient cells suggesting that the bias originated from preferential accurate Pol η bypass of 8-oxoG on the lagging strand instead of any impact of MMR. Thus, human MMR appears to have either a limited role in removing dA paired with 8-oxoG or functions relatively equally between the leading and lagging strands. The underlying reason for HMCES to preferentially limit 8-oxoG mutagenesis on the leading strand is currently unclear, especially considering the synthetic lethal phenotype between HMCES deficiency and APOBEC3A expression^50,72^, which damages the lagging strand template^73^.

In conclusion, this study outlines the multi-dimensional mutational landscape of exogenously and endogenously induced oxidative damage and the consequences of topology on this landscape while providing mechanistic insight into primary, secondary, and tertiary strategies to limit 8-oxoG mutagenesis in human cells. Variants and loss of multiple of the factors in this study lead to cancer and neurodegenerative disease including OGG1 and Pol η. The robust bioinformatics pipeline and exhaustive topological analysis can also be used as a blueprint and foundation to develop a database of holistic multi-dimensional mutational signatures to explore mechanism and drug targets. Future research should explore topological mutagenesis studies of various types of damage and genetic conditions in human cells. This research should also aim to understand the interplay between these pathways and identify potential therapeutic targets for future interventions.

## Materials and Methods

### Data Availability

Full mutation lists used for analysis are available in Supplemental Table 2. hTERT-RPE-1 *POLH*^−/−^ VCF files were available from^47^ available on Mendeley Data server (https://doi.org/10.17632/jkjkpvgxyd.1). MUTYH^−/−^ and OGG1^−/−^ VCF files were accessed from the supplementary dataset S01 from^17^. CLAPS-seq 8-oxoG lesion mapping data from^36^ was obtained from Gene Expression Omnibus (GEO) (https://www.ncbi.nlm.nih.gov/geo/) under accession number GSE181312. Publicly available lists of tumor mutations were obtained from the International Cancer Genome Consortium (ICGC) at https://dcc.icgc.org/api/v1/download?fn=/PCAWG/consensus_snv_indel/final_consensus_passonly.snv_mnv_indel.icgc.public.maf.gz. All custom scripts for mutation and lesion analyses are available at the S-RobertsLab GitHub (https://github.com/S-RobertsLab/Cordero-et-al.-2024).

### System Information

All computational analyses were performed on Linux, specifically Ubuntu 22.04.03 LTS. Data analysis was conducted using Python v3.10.12, Python v2.7.18, Perl v5.34.0 and R v4.1.2 (unless a virtual environment was required). Further system, software, library versions, and hardware information is available on request.

### Cell Culture

hTERT-RPE-1 p53^−/−^ cells were cultured in DMEM (Thermo Fisher Cat No. 11965092 supplemented with 7.5% FBS, 1X Glutamax (Thermo Fisher Cat No. 35050061), 1X Non-Essential Amino Acids (Thermo Fisher Cat No. 11140035, and 1X Penicillin-Streptomycin (Thermo Fisher Cat No. 15140122) at 37° C in 5% CO2. Cells were single-cell cloned with cloning rings. Single-cell clone parental cells were transfected with pSpCas9(BB)-2A-Puro 2 (Addgene Cat No. 48139) that contain guide RNAs that target the intron-exon junction of the second exon of HMCES (5’-TTGCGCCTACCAGGATCGGC and 5’-ACTTTAGACGGTGGTCACGG). Cells were selected with 15 μg/mL puromycin for two days prior to plating for individual clones because hTERT-RPE-1 p53^−/−^ are already mildly puromycin resistant. Clones were screened by immunoblotting for loss of HMCES expression with antibodies raised against the middle and C-terminus of the protein and were hypersensitive to KBrO_3_ as previously described^48^.

### Long-term Mutagenesis Assay

For long-term mutagenesis assays, pooled parental and a single-cell clone (Clone 3) of WT hTERT-RPE-1 p53^−/−^ (generously provided by Daniel Durocher, University of Toronto) as well as three individual single-cell HMCES^−/−^ clones of HMCES knockouts (Clone3.1, Clone 3.3, and Clone 3.4) were seeded into 10cm dishes and carried continuously in the presence or absence of 250µM KBrO_3_ for 100 generations (3 months). Each passage, cells were seeded at similar cell numbers (20% confluency) and carried until 80% confluent at which point they were passaged again. After 100 generations (24 passages), each cell line was single-cell cloned and two of each clone (WT pool, WT Clone 3, HMCES^−/−^ Clones 3.1, 3.3, 3.4) were harvested for genomic DNA (Promega Cat No. A1120). Genomic DNA was submitted for 150 bp paired-end Illumina dep-sequencing sequencing targeting 30X depth at Vanderbilt University’s VANTAGE Next Generation-Sequencing core.

### Sequencing Alignment

Results were aligned to the Genome Reference Consortium Human Build 37 (GRCh37/hg19) using the Burrows-Wheeler Aligner (BWA) `mem` algorithm on default parameters (BWA v0.7.17). The resulting Sequence Alignment/Map (SAM) files, which contain aligned sequence reads, were compressed into Binary Alignment/Map (BAM) format using `samtools` `view` (samtools v1.13 using htslib v1.13+ds). [Note: All `samtools` steps were run using default parameters to maintain a standard approach] After compression, the BAM files were sorted based on genomic coordinates using `samtools` `sort` to prepare for removal of duplicate reads which can arise from PCR amplification artifacts during sequencing. These were removed using `samtools` `rmdup` so it would not have an impact on downstream variant calling and analysis. These final BAM files were converted to MPILEUP files using `samtools` `mpileup`. The final BAM and MPILEUP files were used to call mutations from multiple mutation callers.

### Mutation Calling

The BAM files were processed with Strelka2 (v2.9.10), Manta (v1.6.0), and Somatic Sniper (v1.0.5.0) while the MPILEUP files were processed using VarScan2 (v2.3). SNVs and INDELs were called using `VarScan` `somatic` comparing treated cells to untreated counterparts with the following parameters changed `-min-coverage 10 –min-var-freq 0.2 –somatic-p-value 0.05 –min-freq-for-hom 0.9 –min-avg-qual 30` to reduce artifacts of mutation calling. The resulting SNV and INDEL files were split into germline, somatic, and loss of heterozygosity (LOH) files using `VarScan` `processSomatic` on default parameters, to split the results and isolate the high confidence somatic SNV and INDEL mutation calls which were used to identify consensus mutations.

The BAM files were initially compared to their corresponding normal counterparts utilizing Manta’s structural variant pipeline^74^, employing default parameters to detect small INDEL candidates for input into Strelka2. Strelka2 was run on default parameters comparing tumor to normal using hg19 and Manta’s INDEL candidates for the tumor/normal pair. The resulting SNV and INDEL mutation calls were used to identify consensus mutations. The BAM files were also used to create a third set of SNV calls using Somatic Sniper on default parameters except `-Q 40 –G –L` which requires a minimum somatic score of 40 as recommended by the developers for BWA aligned reads, and not report loss of heterozygosity (LOH) and gain of reference (GOR) mutations in the final output to reduce the likelihood of false positives. The resulting SNV mutation calls were also used to identify consensus mutations.

To account for artifacts of mutation calling and sequencing from different callers, we took the consensus from all three callers (Strelka2, Somatic Sniper, and VarScan2) for SNVs and the consensus from both Strelka2 and VarScan2 for INDELs. This was done using a custom Python script requiring mutations to be present in all sets of mutation calls for the sample. Then all the separate consensus mutations were pooled, and mutations present in more than one sample were omitted due to a high likelihood of being a germline mutation or artifact of sequencing and mutation calling. The concatenated mutation calls were then split into separate sets based on treatment, genotype, or both depending on the analysis.

### Processing of 8-oxoG Lesion Data

CLAPS-seq FASTQ files were aligned to hg19 using the `bwa-mem` algorithm on default parameters. The resulting SAM files were processed using a custom Python script to convert the SAM file into a BED file. The script filtered out reads that did not align with a CIGAR score of 150M. It also filtered reads keeping ones that aligned to chromosome 1-22, X, or Y. It took the reads passing this filter and checked the bitwise flag for 0 (complemented) or 16 (reverse complemented) and processed the proper alignment position (either 5’ or 3’ of the top strand) to determine the base pair position where the lesion occurred. We then filtered the custom BED file for positions where there was a G at that context which removed reads which were assumed to be false positives reported by the authors. The resulting BED files were converted to a VCF format using a custom Python script to process this data through other programs, like vcf2maf and the nucleosome profiling.

### Mutation Signature Generation

We generated mutation signatures from cell mutations using SigProfilerExtractor^32^ (v1.1.23) on default parameters with a minimum and maximum of 1 and 5 signatures respectively. The most stable number of signatures was 2 for both SNV and INDEL mutation signatures, which was used for all subsequent comparisons.

### Correlation of INDEL signature with SBS18

The PCAWG data was analyzed using the MutationalPatterns package in R^35^ to conduct non-negative matrix factorization (NMF). This analysis incorporated the COSMIC SBS and INDEL signatures, along with an additional custom INDEL signature derived from the NMF results of KBrO_3_ treated cells. Tumors were deemed positive for SBS18 and the custom INDEL signature if they exhibited a minimum of 20 mutations associated with each signature. These samples were plotted with the log_2_ transformed number of mutations. The Pearson correlation coefficient was computed based on the mutation count per sample for each signature.

### Strand Asymmetry

Replication strand asymmetry was calculated using AsymTools2^53^ on default parameters. AsymTools2 only works with MAF files, so we converted our VCF files to MAF files using vcf2maf (v1.6.21)^75^ (https://github.com/mskcc/vcf2maf) and VEP (v102)^76^ (https://useast.ensembl.org/info/docs/tools/vep/index.html). The results were filtered and combined so that leading strand events were combined and lagging strand events were combined into a singular figure. Transcribed strand asymmetry was calculated using a custom Python script using a similar approach to AsymTools2. The script takes an RPE-1 transcribed gene list from GEO accession number <u>GSE146121</u> and cross-references the list with the UCSC hg19 gene list. This provided us with a gene list that was actively transcribed in RPE-1 cells, which we then compared with mutations and lesions. Mutations mapping to the top strand with a G base were considered to be on the (+) strand and mutations mapping with a C base were considered to be on the (-) strand. By analyzing the gene’s orientation, we were able to ascertain the strand on which the event took place and subsequently compare the occurrences of each event on both the transcribed and non-transcribed strands. To normalize the events, the event counts were divided by the total length of all genes in the list, resulting in the unit of events/Mb. The results were similar to what was represented in the AsymTools2 results, however, were specific to the cell line and had a higher resolution since transcribed regions were not binned but were measured at single-nucleotide resolution.

### Chromatin State & Nucleosome Profiling

Chromatin states were assessed by mapping mutations and lesions to chromatin states from the epithelial cell HMM chromatin maps (https://genome.ucsc.edu/cgi-bin/hgFileUi?db=hg19&g=wgEncodeBroadHmm). To standardize events by Mb in each region target size, we conducted a division of the events by the cumulative region size. Subsequently, we converted the obtained results to events/Mb based on the HMM map. The order of heterochromatin to euchromatin was determined by the map construction.

Mutations and lesions were intersected with strongly positioned nucleosome dyads following the protocol outlined in^77^ in a 1000 base-pair (bp) window. Expected counts were calculated using genomic trinucleotide mutation or lesion frequencies multiplied by the occurrence of those contexts at each position in the dyad map. The observed counts were divided by the expected counts and log_2_ transformed generate the graphs.

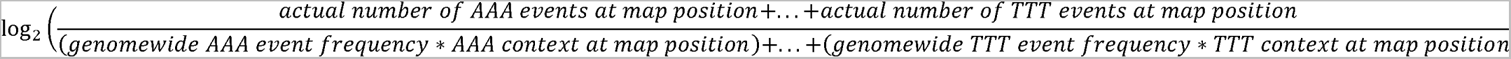

The data was smoothed using a Savitzky–Golay filter with a [200 bp window] with a polynomial order of [3].

### Transcription Factor Profiling

Mutations and lesions were intersected with known active transcription factor binding sites using a map generated from previous work^42^ in a 1000 bp window. Events were counted and graphed using a custom Python script and smoothed using a Savitzky–Golay filter with a 200 bp window with a polynomial order of 3.

### Purification of H. sapiens OGG1 K249Q

A pGEX6P1 vector (N-terminal GST tag) with the *H. sapiens* OGG1 gene bearing the K249Q mutation was obtained from GenScript. For protein expression, the pGEX6P1-OGG1-K249Q vector was transformed into BL21-CondonPlus (DE3) RIPL cells (Agilent). The transformed cells were grown in 2x YT media at 37 °C until an OD_600_ of 0.8 and protein expression induced with 0.5 mM IPTG overnight at 18 °C. The cells were harvested by centrifugation and resuspended in a buffer containing 50 mM HEPES (pH-7.5), 150 mM NaCl, 1 mM DTT, and a protease inhibitor cocktail (Benzamidine, Leupeptin, AEBSF, Pepstatin A). The resuspended cells were lysed by sonication and the lysate clarified by centrifugation. The clarified lysate was loaded onto a GSTrap HP column (Cytiva) equilibrated with 50 mM HEPES (pH-7.5), 150 mM NaCl, and 1 mM DTT, and the protein was eluted in a buffer containing 50 mM HEPES (pH-7.5), 150 mM NaCl, 1 mM DTT, and 50 mM reduced glutathione. Fractions containing GST-OGG1 were loaded onto a Resource S cation exchange column (Cytiva) equilibrated with 50 mM HEPES (pH-6.8), 50 mM NaCl, 1 mM DTT, and 1 mM EDTA, and eluted in a high salt buffer containing 50 mM HEPES (pH-6.8), 1 M NaCl, 1 mM DTT, and 1 mM EDTA. OGG1 was then liberated from the GST-tag by incubation with PreScission Protease for 4 hours in a buffer containing 50 mM HEPES (pH-7.5), 150 mM NaCl, and 1 mM DTT. The cleaved OGG1 protein was rerun over a Resource S cation exchange column (Cytiva), and the eluted protein loaded on a Sephacryl S-200 HR (Cytiva) equilibrated with 50 mM HEPES (pH-7.5), 150 mM NaCl, and 1 mM TCEP. The purified OGG1 fractions were combined, concentrated to 10 mg ml^-1^, and stored at –80 °C.

### Preparation of oligonucleotides

DNA oligonucleotides (oligos) containing 8-oxoG were obtained from TriLink BioTechnologies, and non-damaged oligos were obtained from Integrated DNA Technologies. Each oligo was resuspended at 1 mM in a buffer containing 10 mM Tris (pH-8.0) and 1 mM EDTA. Complimentary oligos (see Supplemental Table 3) were mixed at a 1:1 ratio and annealed by heating to 90 °C followed by a stepwise cooling to 4 °C using a linear gradient at –1 °C min^-1^. The annealed oligos were stored long-term at –20 °C.

### Purification of recombinant human histones

The genes encoding *H. sapien* histones H2A, H2B, H3.2 (C110A), and H4 were cloned into a pet3a expression vector. For histone H2A, H2B, and H3.2 expression, vectors were transformed into T7 Express lysY competent cells (New England Biolabs). For histone H4 expression, the vector was transformed into BL21-CodonPlus (Agilent) competent cells. The cells were grown in minimal media at 37 °C until an OD_600_ of 0.4 was reached, and protein expression induced with 0.4 mM IPTG (H2A, H2B, and H3.2) or 0.3 mM IPTG (H4) for 3-4 hours at 37 °C. The cells were harvested by centrifugation and resuspended in a buffer containing 50 mM Tris (pH-7.5) 100 mM NaCl, 1 mM benzamidine, 1 mM DTT, and 1 mM EDTA. The histones were purified under denaturing conditions using an established method^78,79^. In brief, the resuspended cells were lysed by sonification, inclusion bodies isolated by centrifugation, and the histones extracted from the inclusion bodies under denaturing conditions (6 M Guanidinium chloride). After extraction, the histones were purified using subtractive anion-exchange chromatography and cation-exchange chromatography using gravity flow columns. The purified histones were then dialyzed into H_2_O, lyophilized, and stored at –20 °C.

### Preparation of H2A/H2B Dimers and H3/H4 Tetramers

H2A/H2B dimers and H3/H4 tetramers were prepared using an established method^78,79^. In brief, each individual histone was resuspended in a buffer containing 20 mM Tris (pH-7.5), 6 M guanidinium chloride, and 10 mM DTT. For H2A/H2B dimers, H2A and H2B were mixed at a 1:1 ratio and dialyzed three times against a buffer containing 20 mM Tris (pH-7.5), 2 M NaCl, and 1 mM EDTA. For H3/H4 tetramers, H3 and H4 were mixed at a 1:1 ratio and dialyzed three times against a buffer containing 20 mM Tris (pH-7.5), 2 M NaCl, and 1 mM EDTA. The H2A/H2B dimers and H3/H4 tetramers were subsequently purified over a Sephacryl S-200 HR (Cytiva) in a buffer containing 20 mM Tris (pH-7.5), 2 M NaCl, and 1 mM EDTA. The purified H2A/H2B dimers and H3/H4 tetramers were stored in 50% glycerol at –20 °C.

### Nucleosome assembly and purification

Recombinant nucleosomes were assembled by an established salt-dialysis method^78,79^. In brief, H2A/H2B dimers and H3/H4 tetramers were mixed with DNA in a 2:1:1 molar ratio, respectively, in a buffer containing 20 mM Tris (pH 7.5), 2 M NaCl, and 1 mM EDTA. Stepwise nucleosome assembly was then performed by decreasing the amount of NaCl from 2.0 M NaCl to 1.5 M NaCl, 1.0 NaCl, 0.66 M NaCl, 0.5 M NaCl, 0.25 M NaCl, 0.125 M, and 0 M NaCl over a period of 24 – 26 hours. The reconstituted nucleosomes were heat shocked at 37 °C for 15 minutes to generate uniform DNA positioning and purified by ultracentrifugation over a 10% – 40% sucrose gradient. Final nucleosome purity was determined using native polyacrylamide gel electrophoresis (5%, 59:1 acrylamide:bis-acrylamide), and the purified nucleosomes were stored at 4 °C.

### Cryo-EM sample and grid preparation

For cryo-EM sample preparation, 8oxoG-NCP (5 μM) was mixed with OGG1 K249Q (7.5 μM – 10 μM) in a buffer containing 25 mM HEPES (pH-7.1), 25 mM NaCl, 1 mM TCEP, and 1 mM EDTA. The OGG1-8oxoG-NCP complexes were then incubated at 4 °C for 10 minutes and fixed with glutaraldehyde (0.1%) for 20 minutes. The samples were loaded onto a Superdex S200 Increase 10/300 GL (Cytiva) equilibrated with a buffer containing 50 mM HEPES (pH-7.1), 100 mM NaCl, 1 mM TCEP, and 1 mM EDTA. Fractions containing OGG1-NCP were identified via native polyacrylamide gel electrophoresis (5%, 59:1 acrylamide:bis-acrylamide). The fractions containing the OGG1-8oxoG-NCP complex were then combined and concentrated to 1.5 μM for short-term storage. Gels corresponding to the 8oxoG-NCP_−6_ and 8oxoG-NCP_+4_ samples used for cryo-EM grid preparation can be found in Supplemental Figure 4A and 8A. The samples (3 μL, 1.5 μM) were then applied to a Quantifoil R2/2 300 mesh copper cryo-EM grid at 8 °C and 95% humidity, and the grids plunge frozen in liquid ethane using a Vitrobot Mark IV (Thermo Fisher).

### Cryo-EM Data collection and processing

All cryo-EM data collections were performed on a Titan Krios G3i equipped with Gatan K3 direct electron detector and BioContinuum energy filter at the University of Chicago Advanced Electron Microscopy Core Facility (RRID:SCR_019198). All cryo-EM datasets were processed with cryoSPARC^80^ using the workflows outlined in Supplemental Figure 4 and 8. In brief, the micrographs were corrected for beam-induced drift using Patch Motion Correction and contrast transfer function (CTF) fit using Patch CTF Estimation. The micrographs were then manually curated to exclude micrographs of poor quality. Following micrograph curation, a subset of micrographs was subjected to blob picking to generate initial templates, which were then used for automated template picking. The particle stacks were then extracted from the micrographs and multiple rounds of 2D classification performed. Ab-initio models were then generated using the final particle stacks and several rounds of heterogeneous refinement performed to initially separate 8oxoG-NCP and OGG1-8oxoG-NCP maps.

To improve the interpretability of the 8oxoG-NCP maps, additional 3D-classification was performed using a focus mask for the entry/exit site nucleosomal DNA, which is prone to partially unwrapping from the histone octamer. Following 3D classification, the final particle stacks for each 8oxoG-NCP structure were re-extracted to full box size (600 pixels), and the re-extracted particles subjected to local CTF refinement and non-uniform refinement. The final 8oxoG-NCP maps were then subjected to a B-factor sharpening using PHENIX autosharpen. The final 8oxoG-NCP maps were deposited into the electron microscopy data bank under accession numbers EMD-43595 for 8oxoG-NCP_−6_ and EMD-43600 for 8oxoG-NCP_+4_.

To improve interpretability of the OGG1-8oxoG-NCP maps, 3D-classification was performed using a focus mask for OGG1 and the surrounding nucleosomal DNA. Following 3D-classification, the final particle stacks for each OGG1-8oxoG-NCP structure were re-extracted to full box size (600 pixels), and the re-extracted particles subjected to local CTF refinement and non-uniform refinement. To further improve interpretability of the maps, local refinement (without particle subtraction) was performed using a focus mask for OGG1 and the surrounding nucleosomal DNA or a focus mask for the NCP. A composite map for the OGG1-8oxoG-NCP_−6_ structure was then generated by combining the maps from a non-uniform refinement and two local refinement (OGG1/DNA and NCP local refine) jobs using PHENIX combine focused maps. A composite map for the OGG1-8oxoG-NCP_+4_ structure was then generated by combining the maps from the non-uniform refinement and local refinement (OGG1/DNA local refine) jobs using PHENIX combine focused maps. The final maps were deposited into the electron microscopy data bank under accession numbers EMD-43600 for OGG1-8oxoG-NCP_−6_ (composite), EMD-43597 for OGG1-8oxoG-NCP_−6_ (consensus), EMD-43598 for OGG1-8oxoG-NCP_−6_ (NCP local refine), EMD-43599 for OGG1-8oxoG-NCP_−6_ (OGG1/DNA local refine), EMD-43601 for OGG1-8oxoG-NCP_+4_ (composite), EMD-43602 for OGG1-8oxoG-NCP_+4_ (consensus), and EMD-43603 for OGG1-8oxoG-NCP_+4_ (OGG1/DNA local refine).

### Model building and refinement

All model building and refinement was performed iteratively using University of California San Francisco (UCSF) Chimera^81^, PHENIX^82^, and COOT^83^. An initial nucleosome model was generated using a previously determined cryo-EM structure of a nucleosome containing an AP-site (PDB: 7U52)^63^. The initial OGG1 model was generated from a previously determined X-ray crystal structure of an OGG1-8oxoG-DNA complex (PDB:1EBM)^45^. The models for each respective structure were rigid body docked into the cryo-EM map using UCSF Chimera^81^. The models were then refined in PHENIX^82^ using protein and nucleic acid secondary structure restraints, and manual adjustments to the models made in COOT^83^. All final models were validated using MolProbity^84^, and model coordinates for each structure were deposited in the Protein Data Bank (PDB) under accession numbers 8VWS for 8oxoG-NCP_−6_, 8VWT for OGG1-8oxoG-NCP_−6_, 8VWU for 8oxoG-NCP_+4_, 8VWV for OGG1-8oxoG-NCP_+4_.

## Figure Legends

**Supplemental Table 1: Cryo-EM data collection and processing statistics**.

**Supplemental Table 2: List of mutations from whole genome sequencing of WT and HMCES-deficient RPE-1 cells.**

**Supplemental Table 3: Oligonucleotides used for cryo-EM structures of OGG1 bound to 8-oxoG lesions in nucleosomes.**

**Supplemental Figure 1: Non-negative matrix factorization (NMF) mutation signature generation from KBrO_3_ treated and untreated cells.** The two signatures **(A)** were generated through NMF and SBS96A was present only in the treated cells **(B)** suggesting SBS96B is background. The COSMIC breakdown of the KBrO_3_ treated cell signature and percentage attribution **(C)** led to a cosine similarity of 0.986.

**Supplemental Figure 2: Correlation of KBrO_3_-induced INDEL signature with COSMIC SBS Signatures**. **A,** Fractional breakdown of each nucleotide in positions relative to the deleted T base. The overrepresentation of T on the right and not left is due to the convention of mutation callers left aligning the deleted base in homopolymers. **B,** Visualization of INDEL signature and SBS18 correlation as a red line with samples represented by circles.

**Supplemental Figure 3: NMF Analysis on PCAWG tumors filtered for a minimum of 25% SBS18 mutation contribution in their spectra.** The mutation signature output from the INDEL NMF results with associated percentage makeup of the INDEL spectra.

**Supplemental Figure 4: Single particle analysis of OGG1-8oxoG-NCP_−6_**. **A,** Native PAGE gel of the OGG1-8oxoG-NCP_−6_ cryo-EM sample (S) and a 100 bp DNA ladder (L). The nucleosome complex was visualized with ethidium bromide staining. **B,** Flowchart of the data processing pipeline for the OGG1-8oxoG-NCP_−6_ cryo-EM dataset. A representative micrograph (n=3,899) and representative 2D classes from the OGG1-8oxoG-NCP_−6_ cryo-EM dataset are shown. All maps chosen for the downstream processing are labeled by a dotted red box. The final maps, models, and quality assessment for the OGG1-8oxoG-NCP_−6_ and 8oxoG-NCP_−6_ structures can be found in Supplementary Figure 4 and 5, respectively.

**Supplemental Figure 5: OGG1-8oxoG-NCP_−6_ map and model quality assessment A,** Gold-standard Fourier shell correlation (GS-FSC) curves for the OGG1-8oxoG-NCP_−6_ (consensus, black line), OGG1-8oxoG-NCP_−6_ (NCP focus, blue line), and OGG1-8oxoG-NCP_−6_ (OGG1/DNA focus, red line) cryo-EM maps. The dashed line corresponds to FSC-0.143. **B,** Angular distribution heatmap for the 8oxoG-NCP_−6_ cryo-EM maps. **C,** Map-to-model FSC curves for the OGG1-8oxoG-NCP_−6_ model and OGG1-8oxoG-NCP_−6_ composite cryo-EM map. **D,** Final 3.3 Å OGG1-8oxoG-NCP_−6_ composite cryo-EM map shown in three different orientations. **E**, Final OGG1-8oxoG-NCP_−6_ model shown in three different orientations. **F,** Local resolution estimation for the OGG1-8oxoG-NCP_−6_ composite cryo-EM map shown in three different orientations. **G,** Representative segmented density for histones H2A, H2B, H3, and H4 in the OGG1-8oxoG-NCP_−6_ composite cryo-EM map. **H,** Representative segmented density for OGG1 and the nucleosomal DNA (SHL_-5.5_ to SHL_-6.5_) in the OGG1-8oxoG-NCP_−6_ cryo-EM map.

**Supplemental Figure 6: 8oxoG-NCP_−6_ map and model quality assessment A,** Gold-standard Fourier Shell Correlation (GS-FSC) for the 8oxoG-NCP_−6_. The dashed line corresponds to FSC-0.143. **B,** Angular distribution heatmap for the 8oxoG-NCP_−6_ cryo-EM map. **C**, Map-to-model FSC curves for the 8oxoG-NCP_−6_ model and 8oxoG-NCP_−6_ cryo-EM map. The dashed line corresponds to FSC-0.5. D, Final 3.1 Å cryo-EM map of 8oxoG-NCP_-6_ shown in two different orientations. **E,** Local resolution estimation for the 8oxoG-NCP_−6_ cryo-EM map shown in two different orientations. **F,** Final 8oxoG-NCP_−6_ model shown in two different views. **G,** Representative segmented density for histones H2A, H2B, H3, and H4 in the 8oxoG-NCP_−6_ cryo-EM map. **H,** Representative segmented density for the nucleosomal DNA (SHL_-5.5_ to SHL_-6.5_) in the 8oxoG-NCP_−6_ cryo-EM map. The black dashed box highlights the 8-oxoG:C base pair at SHL_−6_. **I,** Representative segmented density for the Watson-Crick 8-oxoG:C base pair at SHL_−6_ in the 8oxoG-NCP_−6_ cryo-EM map.

**Supplemental Figure 7: A conserved OGG1 8oxoG recognition mechanism at SHL_+4_ A,** Structural comparison of OGG1 and the nucleosomal DNA in the OGG1-8oxoG-NCP_−6_, OGG1-8oxoG-NCP_+4_, and OGG1-8oxoG-DNA (1EBM) complexes. **B,** A diagram representing the interactions between OGG1 and the nucleosomal DNA in the OGG1-8oxoG-NCP_+4_ complex identified using PLIP^85^. **C,** Focused view of the nucleosomal DNA at SHL_+3.5_ to SHL_+4.5_ showing the extrahelical 8-oxoG at SHL_+4_. The segmented density for the nucleosomal DNA in the OGG1-8oxoG-NCP_+4_ composite cryo-EM map is shown in transparent grey. An inset of the OGG1 active site is shown highlighting key amino acids important for 8-oxoG recognition and excision. **D,** Structural comparison of the nucleosomal DNA (SHL_+3.5_ to SHL_+4.5_) in the OGG1-8oxoG-NCP_+4_ complex and 8oxoG-NCP_+4_. **E,** OGG1-8oxoG-NCP_−6_ cryo-EM model (left) and a structural model of OGG1 bound to a histone proximal 8-oxoG at SHL_−6_ (right), showing significant clashes between OGG1 and the nucleosome when 8-oxoG is adjacent to the histone octamer. **F,** OGG1-8oxoG-NCP_+4_ cryo-EM model (left) and a structural model of OGG1 bound to a histone proximal 8-oxoG at SHL_+4_ (right), showing significant clashes between OGG1 and the nucleosome when 8-oxoG is adjacent to the histone octamer.

**Supplemental Figure 8: Single particle analysis of OGG1-8oxoG-NCP_+4_**. **A,** Native PAGE gel of the OGG1-8oxoG-NCP_+4_ cryo-EM sample (S) and a 100 bp DNA ladder (L). The nucleosome complex was visualized with ethidium bromide staining. **B,** Flowchart of the data processing pipeline for the OGG1-8oxoG-NCP_+4_ cryo-EM dataset. A representative micrograph (n=3,317) and representative 2D classes from the OGG1-8oxoG-NCP_+4_ cryo-EM dataset are shown. All maps chosen for the downstream processing are labeled by a dotted red box. The final maps, models, and quality assessment for the OGG1-8oxoG-NCP_+4_ and 8oxoG-NCP_+4_ structures can be found in Supplementary Figure 7 and 8, respectively.

**Supplemental Figure 9: OGG1-8oxoG-NCP_+4_ map and model quality assessment A,** Gold-standard Fourier shell correlation (GS-FSC) curves for the OGG1-8oxoG-NCP_+4_ (consensus, black line) and OGG1-8oxoG-NCP_+4_ (OGG1/DNA focus, red line) cryo-EM maps. The dashed line corresponds to FSC-0.143. **B,** Angular distribution heatmap for the 8oxoG-NCP_+4_ cryo-EM maps. **C,** Map-to-model FSC curves for the OGG1-8oxoG-NCP_+4_ model and OGG1-8oxoG-NCP_+4_ composite cryo-EM map. **D,** Final 3.6 Å OGG1-8oxoG-NCP_+4_ composite cryo-EM map shown in three different orientations. **E,** Final OGG1-8oxoG-NCP_+4_ model shown in three different orientations. **F,** Local resolution estimation for the OGG1-8oxoG-NCP_+4_ composite cryo-EM map shown in three different orientations. **G,** Representative segmented density for histones H2A, H2B, H3, and H4 in the OGG1-8oxoG-NCP_+4_ composite cryo-EM map. **H,** Representative segmented density for OGG1 and the nucleosomal DNA (SHL_+3.5_ to SHL_+4.5_) in the OGG1-8oxoG-NCP_+4_ cryo-EM map.

**Supplemental Figure 10: 8oxoG-NCP_+4_ map and model quality assessment A,** Gold-standard Fourier Shell Correlation (GS-FSC) for the 8oxoG-NCP_+4_. The dashed line corresponds to FSC-0.143. **B,** Angular distribution heatmap for the 8oxoG-NCP_+4_ cryo-EM map. **C,** Map-to-model FSC curves for the 8oxoG-NCP_+4_ model and 8oxoG-NCP_+4_ cryo-EM map. The dashed line corresponds to FSC-0.5. **D,** Final 3.0 Å cryo-EM map of 8oxoG-NCP_+4_ shown in three different orientations. **E,** Local resolution estimation for the 8oxoG-NCP_+4_ cryo-EM map shown in two different orientations. **F,** Final 8oxoG-NCP_+4_ model shown in two different views. **G,** Representative segmented density for histones H2A, H2B, H3, and H4 in the 8oxoG-NCP_+4_ cryo-EM map. **H,** Representative segmented density for the nucleosomal DNA (SHL_+3_ to SHL_+4.5_) in the 8oxoG-NCP_+4_ cryo-EM map. The black dashed box highlights the 8-oxoG:C base pair at SHL_+4_. **I,** Representative segmented density for the Watson-Crick 8-oxoG:C base pair at SHL_+4_ in the 8oxoG-NCP_+4_ cryo-EM map.

**Supplemental Figure 11: Spontaneous mutation spectra in untreated human cells with 8-oxoG associated gene knockouts**. **A**, Number of SNVs and INDELs per genome in untreated human cells. Circles indicate independent genomes sequenced. Horizontal bars are median values. **B**, Mutation spectra plots of spontaneous mutations in human WT, HMCES^−/−^, POLH^−/−^, MUTYH^−/−^, and OGG1^−/−^ cells along with the mutation spectra from SBS18.

**Supplemental Figure 12: The impact of chromatin state on 8-oxodG mutagenesis in human cells**. Four binned broad chromatin regions are dictated by the ChromHMM map and within groups are sorted from left to right in becoming increasingly more euchromatic. Normalized mapped mutations were graphed for **(A)** HMCES^−/−^ RPE-1 cells, **(B)** POLH^−/−^ RPE-1 cells and **(C)** mutations from PCAWG that have a 70% or greater attribution to SBS18. The mean is displayed as a dashed red line and standard deviations on either side are displayed as a dashed blue line.

**Supplemental Figure 13: The impact of chromatin structure on 8-oxodG mutagenesis in human cells.** The top graphs represent translational periodicity of log2(observed/expected) of mutations between nucleosomes. Nucleosome bound DNA is represented in blue and linker DNA is represented in red. The bottom graphs represent the rotational periodicity of the log2(observed/expected) of mutations within the nucleosome where DNA that is inward histone facing DNA relative to the nucleosome is displayed in gold while outward solvent facing DNA relative to the nucleosome is displayed in purple. A binomial fit of the data is overlayed in a dashed green line. In both figures, actual data points are displayed in gray. This was done for **(A)** HMCES^−/−^ RPE-1 cells, **(B)** POLH^−/−^ RPE-1 cells and **(C)** mutations from PCAWG that have a 70% or greater attribution to SBS18.

