## Supplementary figures and images for "Hierarchical determinants of the oxidation-induced mutational landscape in human cells"

### Supplemental Figure 1

#### Supplemental Figure 1

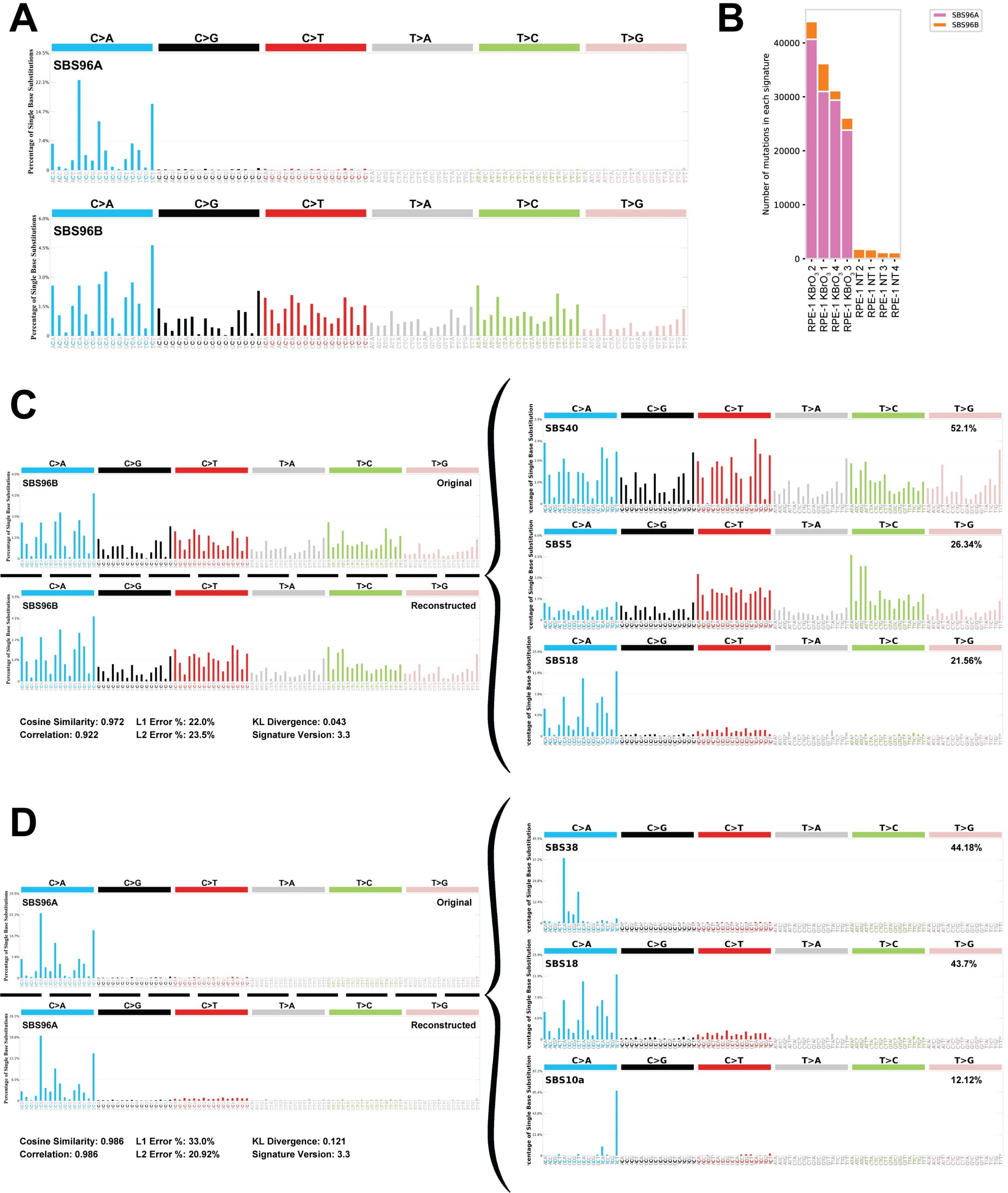

### Supplemental Figure 2

#### Supplemental Figure 2

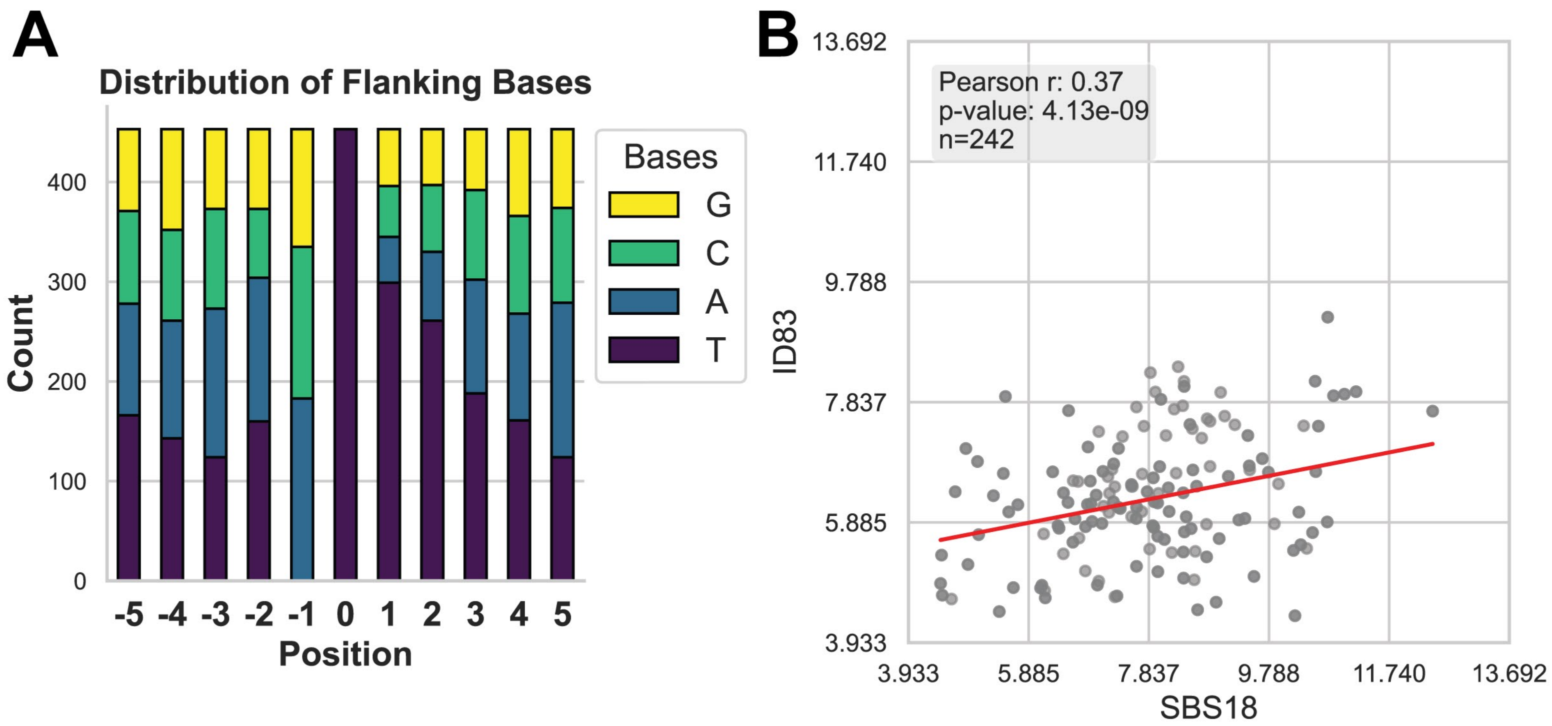

### Supplemental Figure 3

#### Supplemental Figure 3

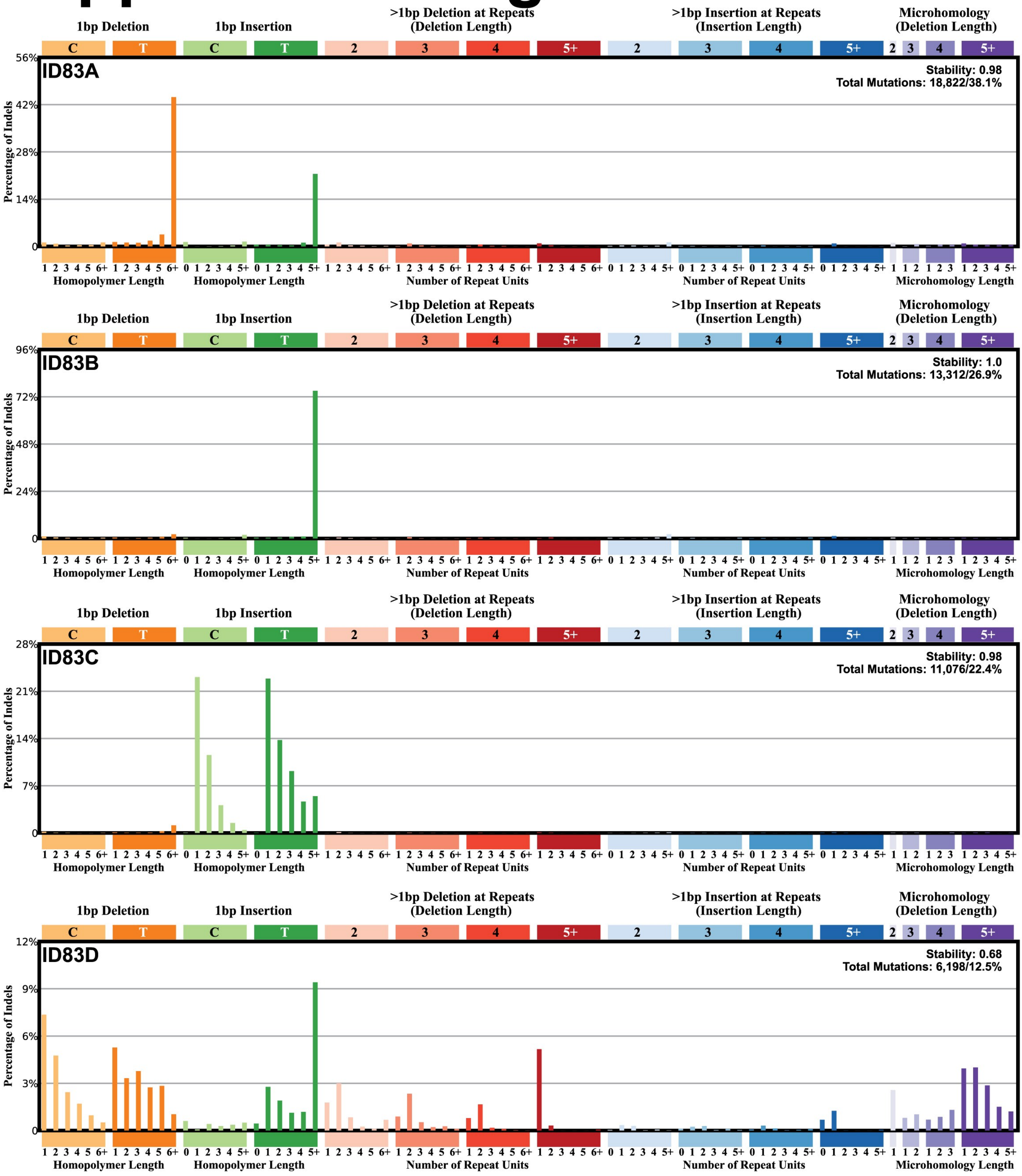

### Supplemental Figure 4

#### Supplemental Figure 4

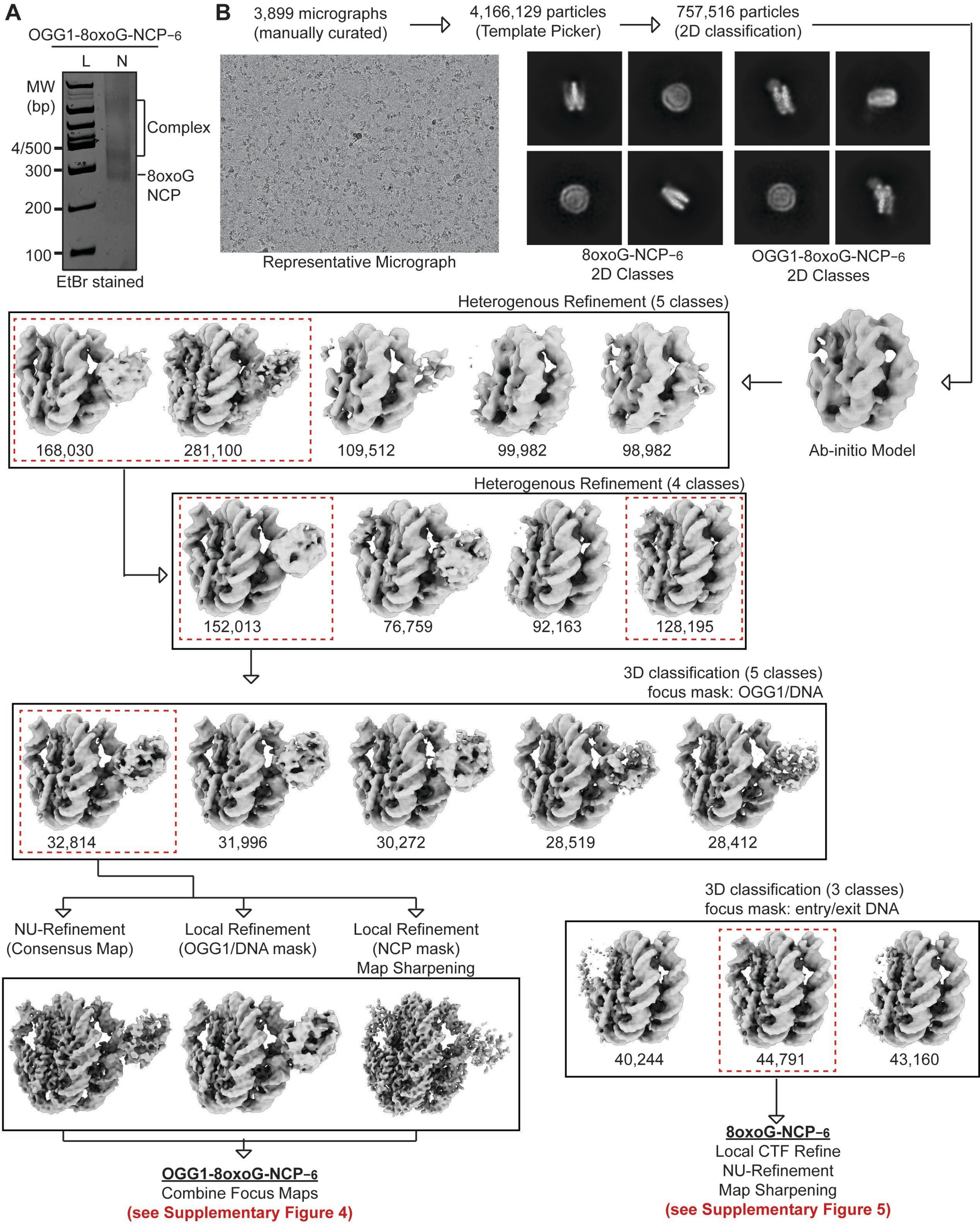

### Supplemental Figure 5

#### Supplemental Figure 5

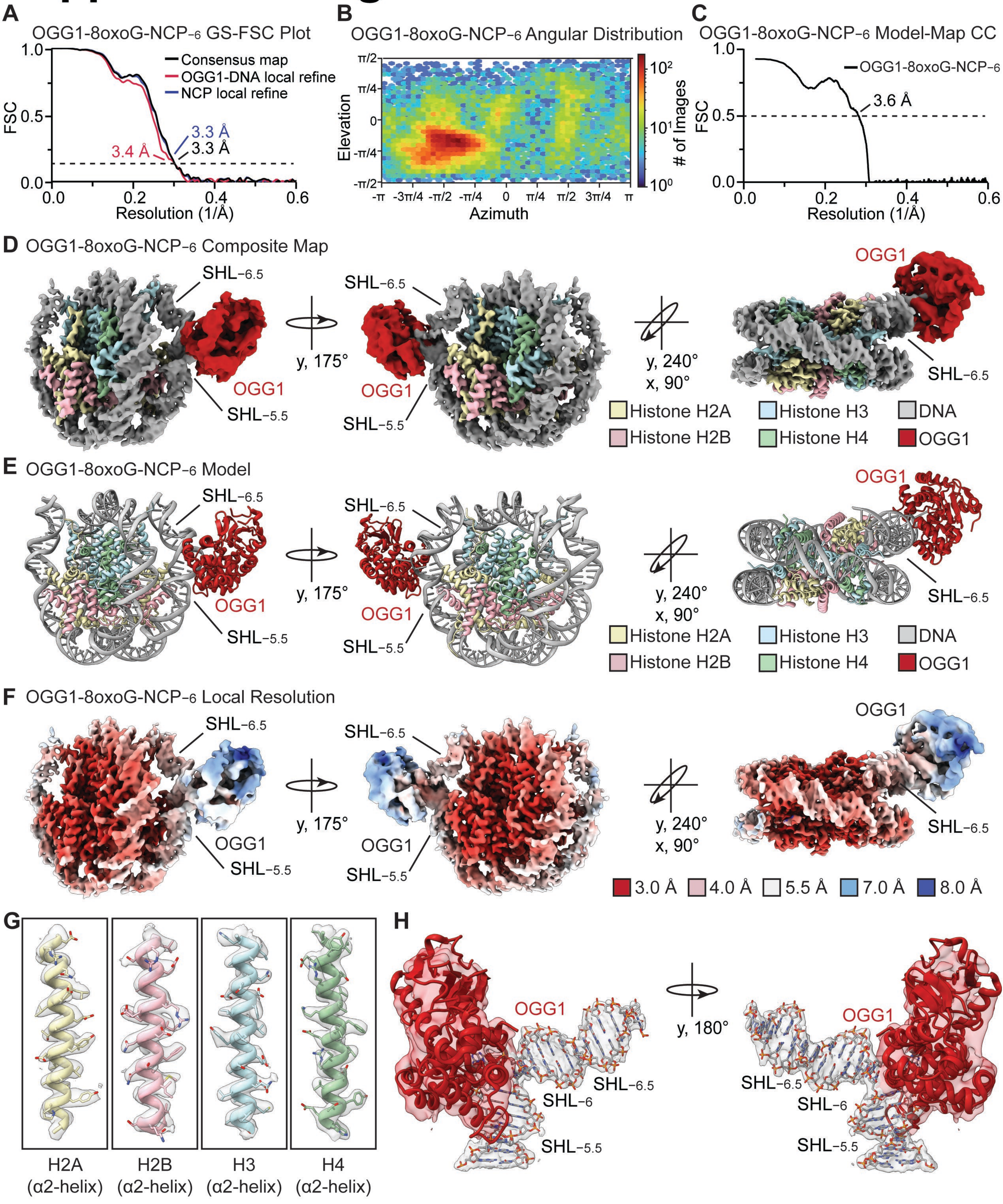

### Supplemental Figure 6

#### Supplemental Figure 6

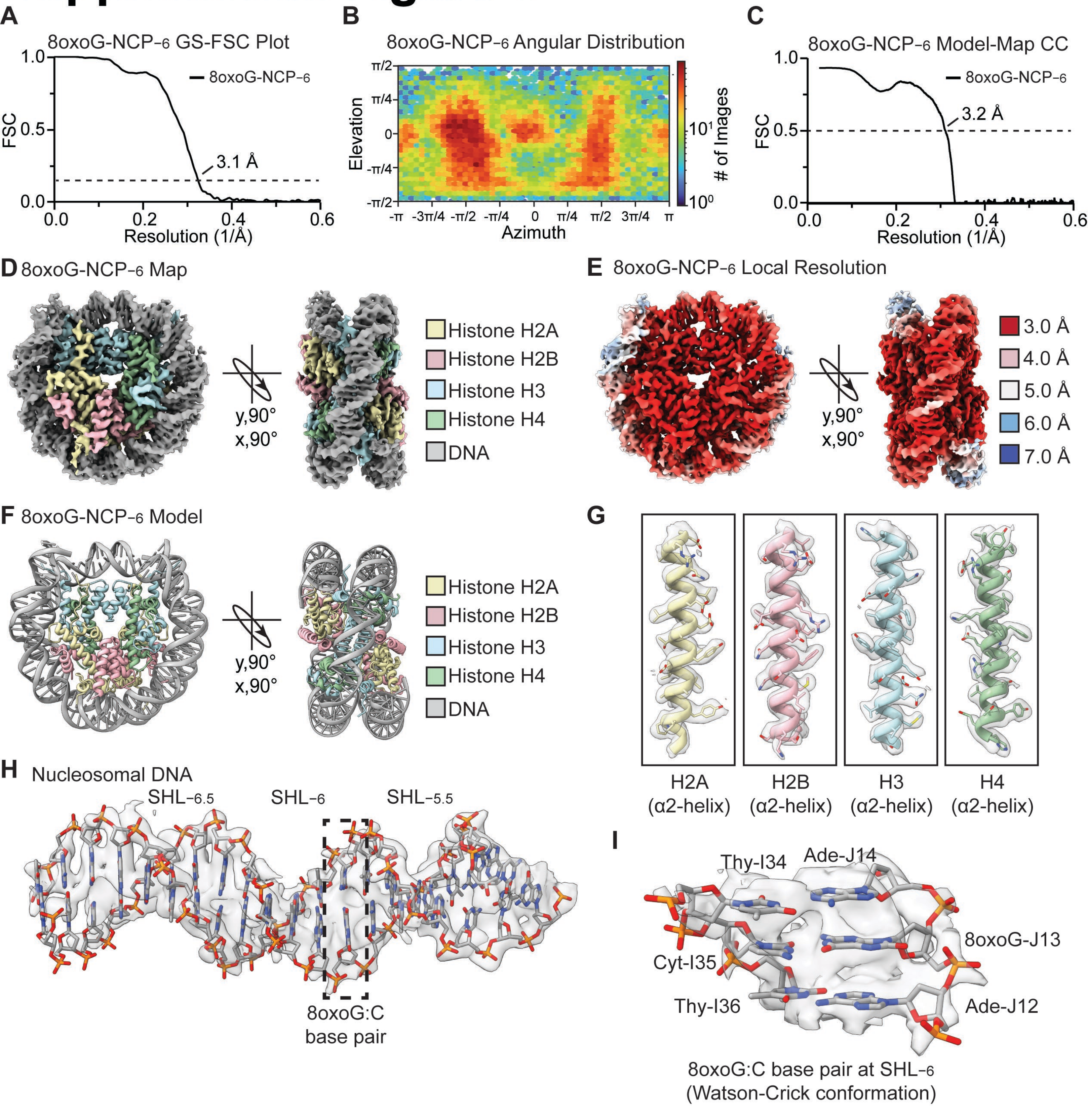

### Supplemental Figure 7

#### Supplemental Figure 7

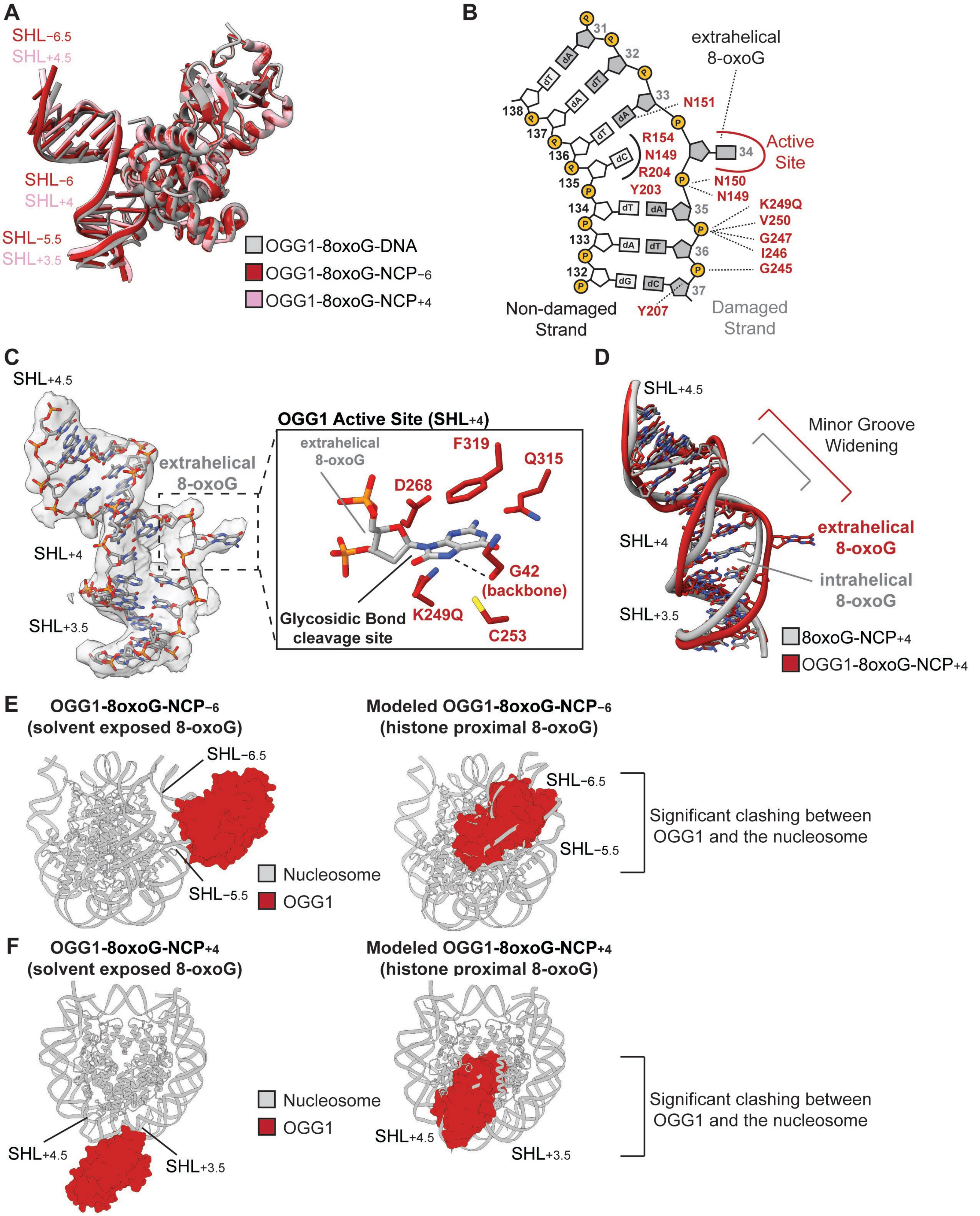

### Supplemental Figure 8

#### Supplemental Figure 8

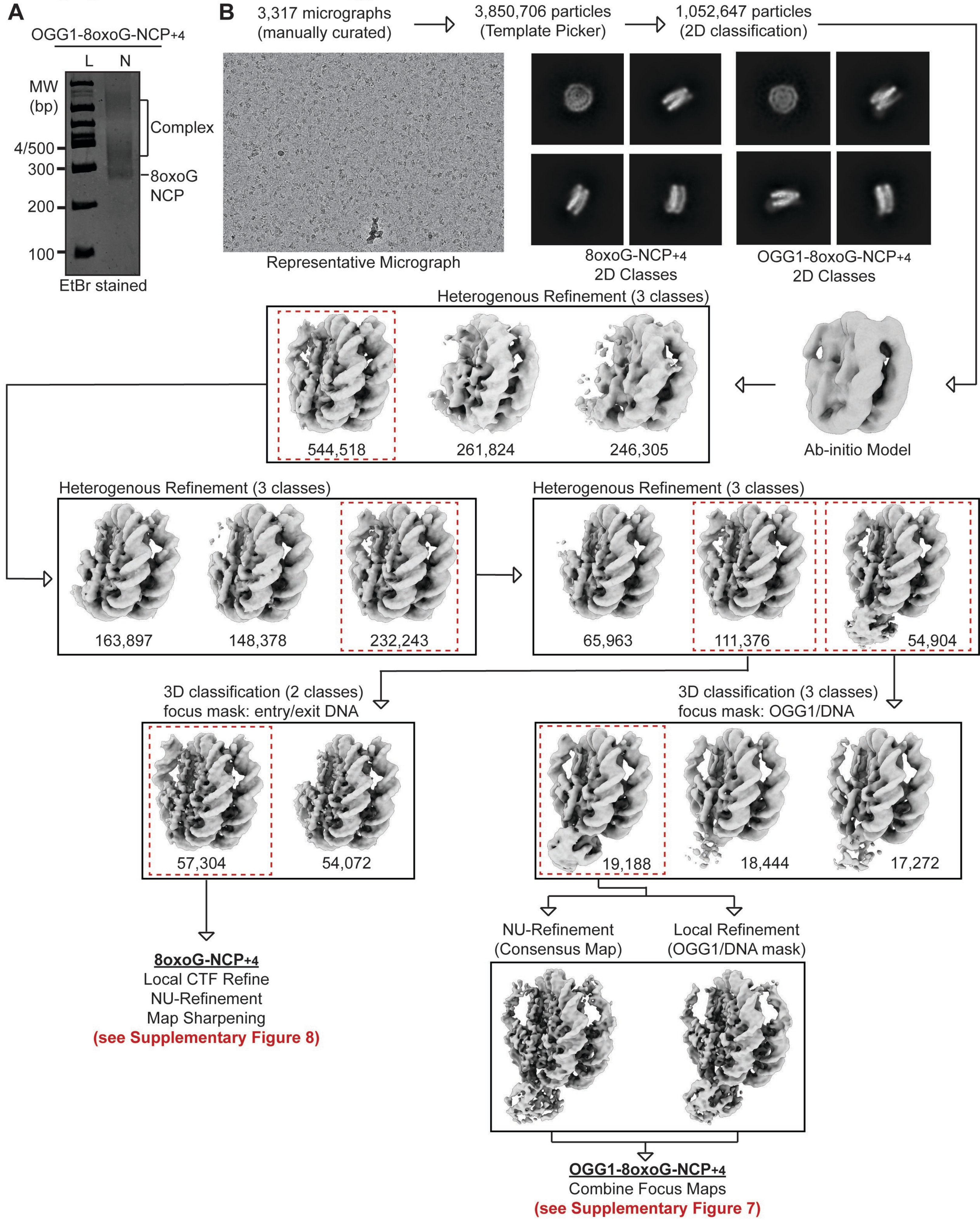

### Supplemental Figure 9

#### Supplemental Figure 9

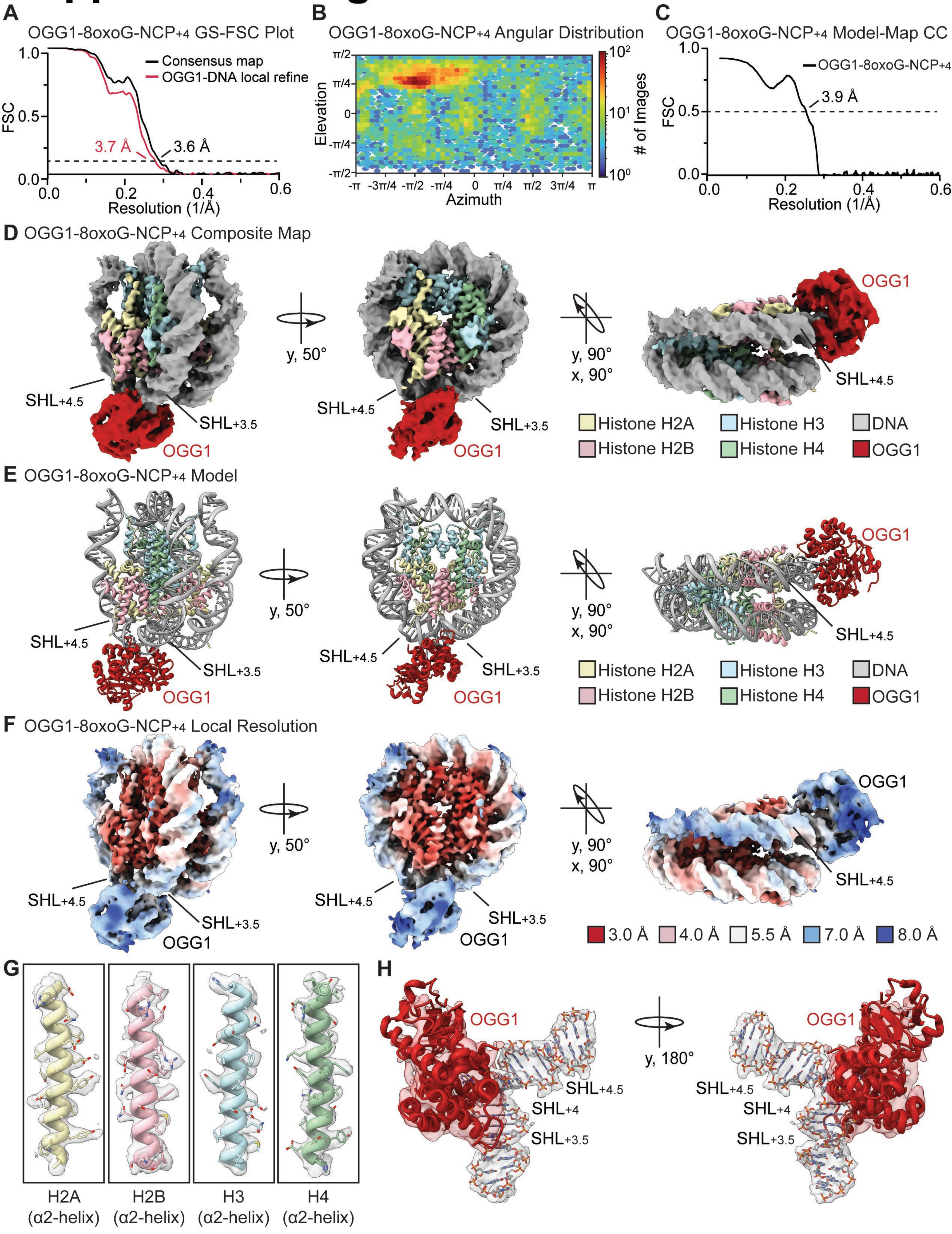

### Supplemental Figure 10

#### Supplemental Figure 10

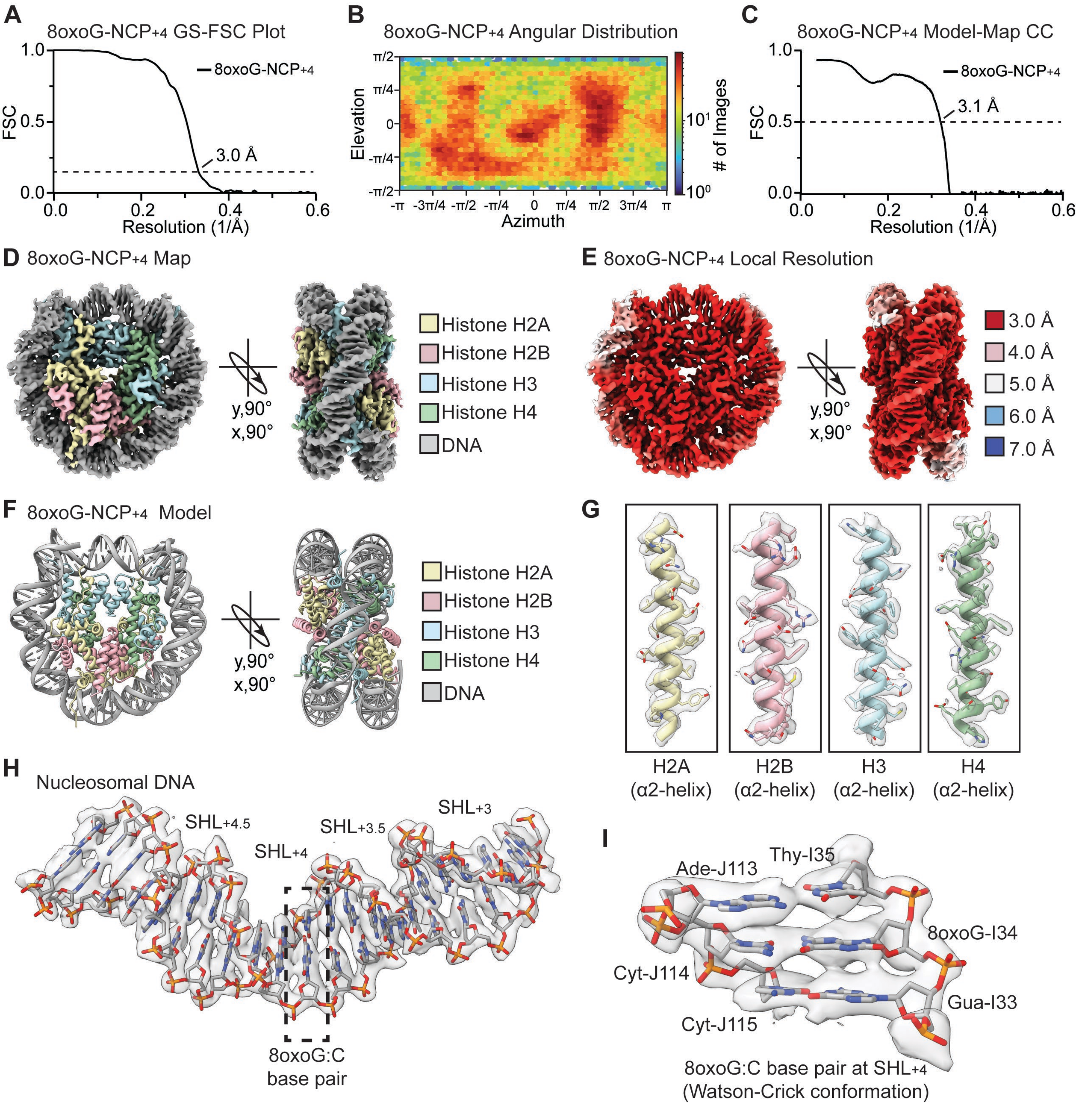

### Supplemental Figure 11

#### Supplemental Figure 11

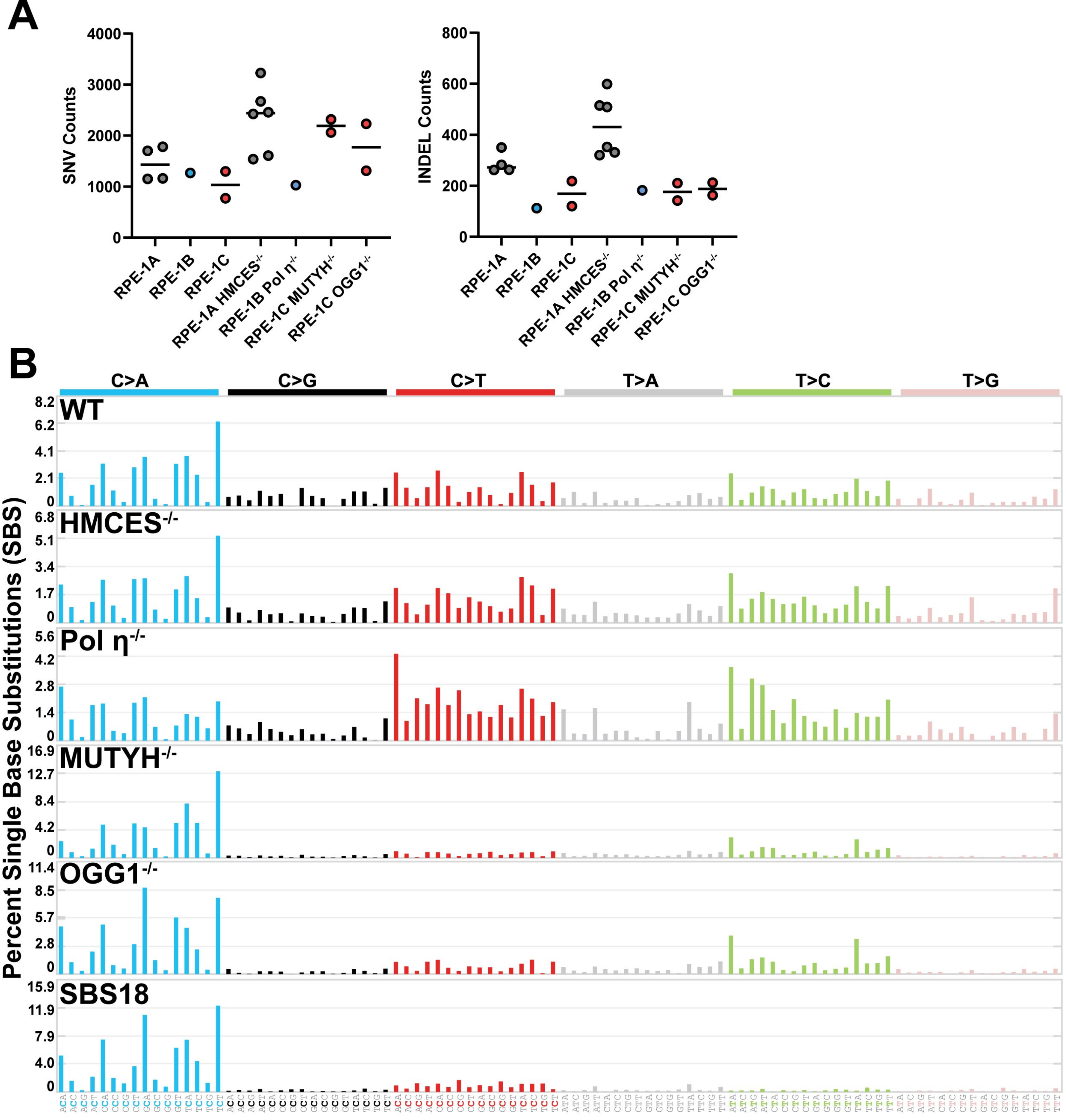

### Supplemental Figure 12

#### Supplemental Figure 12

##### Mutations Across Chromatin Domains

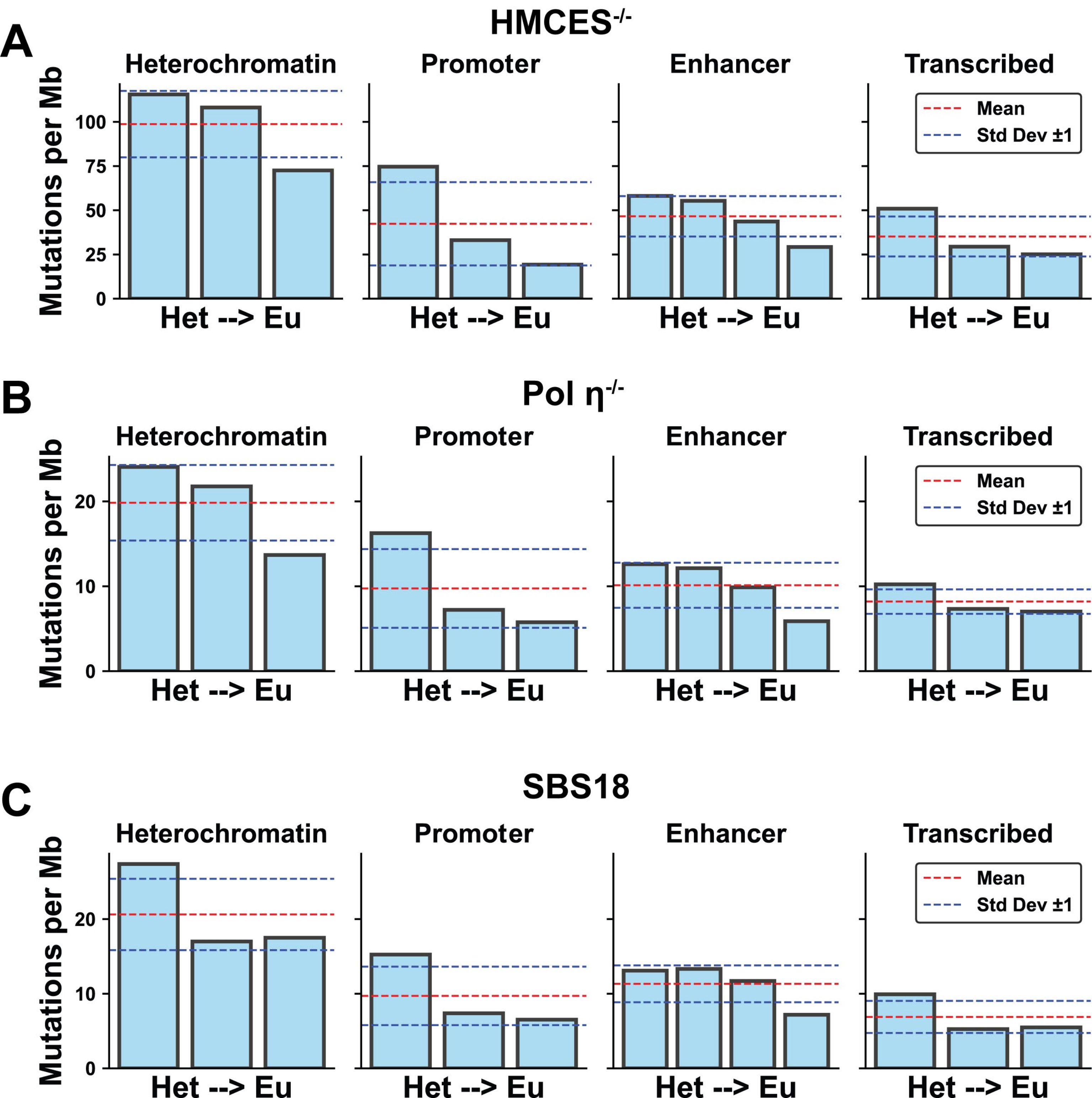

### Supplemental Figure 13

#### Supplemental Figure 13

A

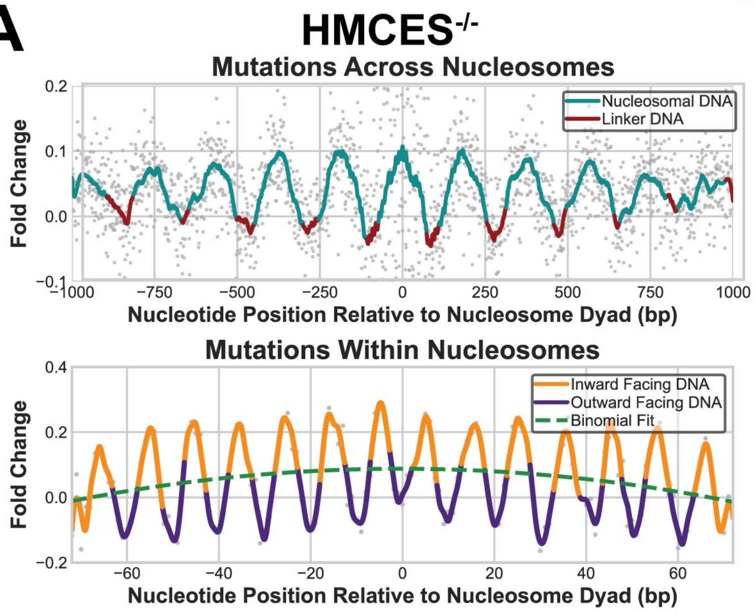


B

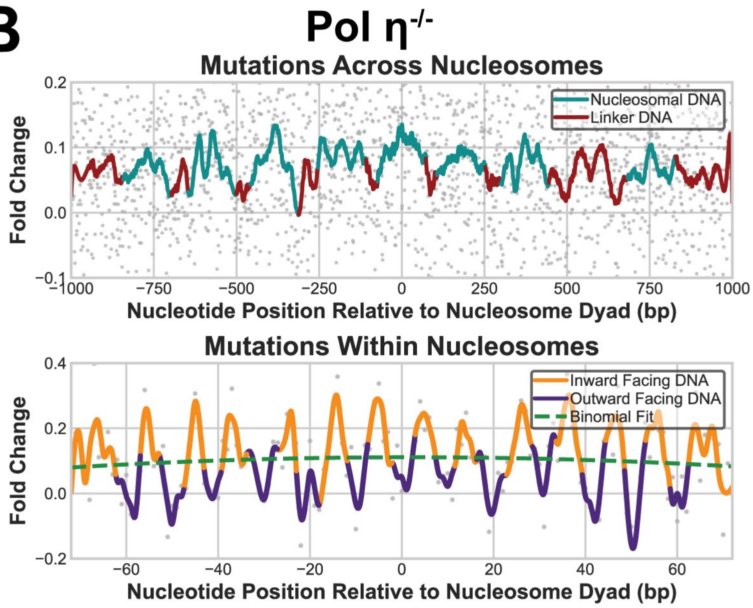


C

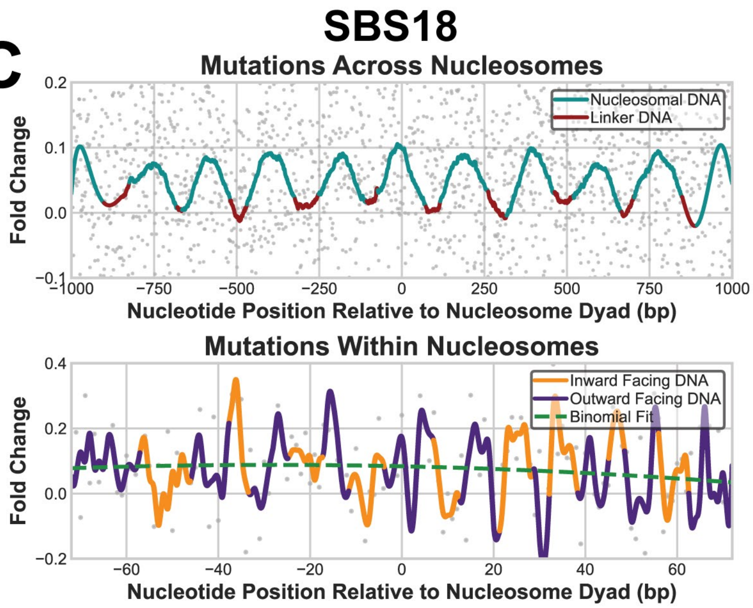
