## Supplemental Table 1 for "Hierarchical determinants of the oxidation-induced mutational landscape in human cells"

| **Data collection and processing** | | | | |
| --- | --- | --- | --- | --- |
| **Dataset** | **OGG1-8oxoG-NCP−6** | | **OGG1-8oxoG-NCP+4** | |
| Magnification | 81,000x | | 81,000x | |
| Voltage (kV) | 300 | | 300 | |
| Electron exposure (e^–^/Å^2^) | 60 | | 60 | |
| Defocus range (μm) | −0.5 to −2.5 | | −0.5 to −2.5 | |
| Pixel size (Å) | 0.534 | | 0.534 | |
| Symmetry imposed | C1 | | C1 | |
| Initial particle number | 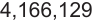 | | 3,850,706 | |
| **Structure** | **8oxoG-NCP−6** | **OGG1-8oxoG-NCP−6** | **8oxoG-NCP+4** | **OGG1-8oxoG-NCP+4** |
| Final particle number | 44,791 | 32,685 | 57,304 | 19,114 |
| Map resolution (Å) | 3.1 | 3.3 | 3.0 | 3.6 |
| FSC threshold | 0.143 | 0.143 | 0.143 | 0.143 |
| PDB accession | 8VWS | 8VWT | 8VWU | 8VWV |
| EMDB accession | EMD-43595 | EMD-43596 | EMD-43600 | EMD-43601 |
| **Refinement** | | | | |
| Initial model used (PDB ID) | 7U52 | 7U52, 1EBM | 7U52 | 7U52, 1EBM |
| Model resolution (Å) | 3.2 | 3.6 | 3.1 | 3.9 |
| FSC threshold | 0.5 | 0.5 | 0.5 | 0.5 |
| **Model composition** | | | | |
| Nonhydrogen atoms | 11,907 | 14,451 | 12021 | 14,474 |
| Protein residues | 743 | 1,061 | 757 | 1,066 |
| Nucleotide | 294 | 294 | 294 | 294 |
| **B factors (Å^2^ )** | | | | |
| Protein | 31.22 | 83.65 | 22.52 | 146.82 |
| Nucleotide | 80.91 | 111.48 | 61.90 | 184.59 |
| **r.m.s. deviations** | | | | |
| Bond Length (Å) (# > 4σ) | 0.004 (0) | 0.005 (2) | 0.003 (0) | 0.005 (0) |
| Bond Angles (^o^) (# > 4σ) | 0.619 (1) | 0.886 (33) | 0.558 (0) | 0.868 (27) |
| **Validation** | | | | |
| MolProbity score | 1.27 | 1.67 | 1.23 | 1.53 |
| Clashscore | 5.10 | 6.92 | 4.53 | 6.15 |
| Poor rotamers (%) | 0.00 | 0.79 | 0.00 | 0.68 |
| **Ramachandran plot** | | | | |
| Favored (%) | 98.21 | 95.87 | 98.65 | 96.85 |
| Allowed (%) | 1.79 | 4.13 | 1.35 | 3.15 |
| Disallowed (%) | 0.00 | 0.00 | 0.00 | 0.00 |
