## Supplemental Table 3 for "Hierarchical determinants of the oxidation-induced mutational landscape in human cells"

| **Oligo** | **Sequence (5′ – 3′)** |
| --- | --- |
| **8oxoG1-NCP oligos** |  |
| 8oxoG-NCP−6  (I-strand) | ATCGAGAATCCCGGTGCCGAGGCCGCTCAATTGGTCGTAGACAGCTCTAGCACCGCTTAAACGCACGTACGCGCTGTCCCCCGCGTTTTAACCGCCAAGGGGATTACT CCC TAGTCT CCAGGCACGTGTCAGATCTATACATCCGAT |
| 8oxoG-NCP−6  (J-strand) | ATCGGATGTATA**[8oxoG]**ATCTGACACGTGCCTGGAGACTAGGGAGTAATCCCCTTGGCGGTTAAAACGCGGGGGACAGCGCGTACGTGCGTTTAAGCGGTGCTAGAGCTGTCTACGACCAATTGAGCGGCCTCGGCACCGGGATTCTCGAT |
| **8oxoG2-NCP oligos** |  |
| 8oxoG-NCP+4  (I-strand) | ATCGAGAATCCCGGTGCCGAGGCCGCTCAATTG**[8oxoG]**TCGTAGACAGCTCTAGCACCGCTTAAACGCACGTACGCGCTGTCCCCCGCGTTTTAACCGCCAAGGGGATTACTCCCTAGTCTCCAGGCACGTGTCAGATATATACATCCGAT |
| 8oxoG-NCP+4  (J-strand) | ATCGGATGTATATATCTGACACGTGCCTGGAGACTAGGGAGTAATCCCCTTGGCGGTTAAAACGCGGGGGACAGCGCGTACGTGCGTTTAAGCGGTGCTAGAGCTGTCTACGACCAATTGAGCGGCCTCGGCACCGGGATTCTCGAT |
